# Protein-Coding Variants Implicate Novel Genes Related to Lipid Homeostasis Contributing to Body Fat Distribution

**DOI:** 10.1101/352674

**Authors:** Anne E Justice, Tugce Karaderi, Heather M Highland, Kristin L Young, Mariaelisa Graff, Yingchang Lu, Valérie Turcot, Paul L Auer, Rebecca S Fine, Xiuqing Guo, Claudia Schurmann, Adelheid Lempradl, Eirini Marouli, Anubha Mahajan, Thomas W Winkler, Adam E Locke, Carolina Medina-Gomez, Tõnu Esko, Sailaja Vedantam, Ayush Giri, Ken Sin Lo, Tamuno Alfred, Poorva Mudgal, Maggie CY Ng, Nancy L Heard-Costa, Mary F Feitosa, Alisa K Manning, Sara M Willems, Suthesh Sivapalaratnam, Goncalo Abecasis, Dewan S Alam, Matthew Allison, Philippe Amouyel, Zorayr Arzumanyan, Beverley Balkau, Lisa Bastarache, Sven Bergmann, Lawrence F Bielak, Matthias Blüher, Michael Boehnke, Heiner Boeing, Eric Boerwinkle, Carsten A Boger, Jette Bork-Jensen, Erwin P Bottinger, Donald W Bowden, Ivan Brandslund, Linda Broer, Amber A Burt, Adam S Butterworth, Mark J Caulfield, Giancarlo Cesana, John C Chambers, Daniel I Chasman, Yii-Der Ida Chen, Rajiv Chowdhury, Cramer Christensen, Audrey Y Chu, Francis S Collins, James P Cook, Amanda J Cox, David S Crosslin, John Danesh, Paul IW de Bakker, Simon de Denus, Renee de Mutsert, George Dedoussis, Ellen W Demerath, Joe G Dennis, Josh C Denny, Emanuele Di Angelantonio, Marcus Dorr, Fotios Drenos, Marie-Pierre Dube, Alison M Dunning, Douglas F Easton, Paul Elliott, Evangelos Evangelou, Aliki-Eleni Farmaki, Shuang Feng, Ele Ferrannini, Jean Ferrieres, Jose C Florez, Myriam Fornage, Caroline S Fox, Paul W Franks, Nele Friedrich, Wei Gan, Ilaria Gandin, Paolo Gasparini, Vilmantas Giedraitis, Giorgia Girotto, Mathias Gorski, Harald Grallert, Niels Grarup, Megan L Grove, Stefan Gustafsson, Jeff Haessler, Torben Hansen, Andrew T Hattersley, Caroline Hayward, Iris M Heid, Oddgeir L Holmen, G Kees Hovingh, Joanna MM Howson, Yao Hu, Yi-Jen Hung, Kristian Hveem, M Arfan Ikram, Erik Ingelsson, Anne U Jackson, Gail P Jarvik, Yucheng Jia, Torben Jørgensen, Pekka Jousilahti, Johanne M Justesen, Bratati Kahali, Maria Karaleftheri, Sharon LR Kardia, Fredrik Karpe, Frank Kee, Hidetoshi Kitajima, Pirjo Komulainen, Jaspal S Kooner, Peter Kovacs, Bernhard K Kramer, Kari Kuulasmaa, Johanna Kuusisto, Markku Laakso, Timo A Lakka, David Lamparter, Leslie A Lange, Claudia Langenberg, Eric B Larson, Nanette R Lee, Wen-Jane Lee, Terho Lehtimäki, Cora E Lewis, Huaixing Li, Jin Li, Ruifang Li-Gao, Li-An Lin, Xu Lin, Lars Lind, Jaana Lindström, Allan Linneberg, Ching-Ti Liu, Dajiang J Liu, Jian’an Luan, Leo-Pekka Lyytikäinen, Stuart MacGregor, Reedik Mägi, Satu Männistö, Gaëlle Marenne, Jonathan Marten, Nicholas GD Masca, Mark I McCarthy, Karina Meidtner, Evelin Mihailov, Leena Moilanen, Marie Moitry, Dennis O Mook-Kanamori, Anna Morgan, Andrew P Morris, Martina Muller-Nurasyid, Patricia B Munroe, Narisu Narisu, Christopher P Nelson, Matt Neville, Ioanna Ntalla, Jeffrey R O’Connel, Katharine R Owen, Oluf Pedersen, Gina M Peloso, Craig E Pennell, Markus Perola, James A Perry, John RB Perry, Tune H Pers, Ailith Pirie, Ozren Polasek, Olli T Raitakari, Asif Rasheed, Chelsea K Raulerson, Rainer Rauramaa, Dermot F Reilly, Alex P Reiner, Paul M Ridker, Manuel A Rivas, Neil R Robertson, Antonietta Robino, Igor Rudan, Katherine S Ruth, Danish Saleheen, Veikko Salomaa, Nilesh J Samani, Pamela J Schreiner, Matthias B Schulze, Robert A Scott, Marcelo P Segura-Lepe, Xueling Sim, Andrew J Slater, Kerrin S Small, Blair H Smith, Jennifer A Smith, Lorraine Southam, Timothy D Spector, Elizabeth K Speliotes, Kari Stefansson, Valgerdur Steinthorsdottir, Kathleen E Stirrups, Konstantin Strauch, Heather M Stringham, Michael Stumvoll, Liang Sun, Praveen Surendran, Karin MA Swart, Jean-Claude Tardif, Kent D Taylor, Alexander Teumer, Deborah J Thompson, Gudmar Thorleifsson, Unnur Thorsteinsdottir, Betina H Thuesen, Anke Tönjes, Mina Torres, Emmanouil Tsafantakis, Jaakko Tuomilehto, André G Uitterlinden, Matti Uusitupa, Cornelia M van Duijn, Mauno Vanhala, Rohit Varma, Sita H Vermeulen, Henrik Vestergaard, Veronique Vitart, Thomas F Vogt, Dragana Ntalla, Lynne E Wagenknecht, Mark Walker, Lars Wallentin, Feijie Wang, Carol A Wang, Shuai Wang, Nicholas J Wareham, Helen R Warren, Dawn M Waterworth, Jennifer Wessel, Harvey D White, Cristen J Willer, James G Wilson, Andrew R Wood, Ying Wu, Hanieh Yaghootkar, Jie Yao, Laura M Yerges-Armstrong, Robin Young, Eleftheria Zeggini, Xiaowei Zhan, Weihua Zhang, Jing Hua Zhao, Wei Zhao, He Zheng, Wei Zhou, M Carola Zillikens, GoT2D Genes Consortium CHD Exome+ Consortium, EPIC-CVD Consortium, ExomeBP Consortium, Global Lipids Genetic Consortium, InterAct, ReproGen Consortium, T2D-Genes Consortium, The MAGIC Investigators, Fernando Rivadeneira, Ingrid B Borecki, John A Pospisilik, Panos Deloukas, Timothy M Frayling, Guillaume Lettre, Karen L Mohlke, Jerome I Rotter, Zoltan Kutalik, Joel N Hirschhorn, L Adrienne Cupples, Ruth JF Loos, Kari E North, Cecilia M Lindgren

## Abstract

Body fat distribution is a heritable risk factor for a range of adverse health consequences, including hyperlipidemia and type 2 diabetes. To identify protein-coding variants associated with body fat distribution, assessed by waist-to-hip ratio adjusted for body mass index, we analyzed 228,985 predicted coding and splice site variants available on exome arrays in up to 344,369 individuals from five major ancestries for discovery and 132,177 independent European-ancestry individuals for validation. We identified 15 common (minor allele frequency, MAF≥5%) and 9 low frequency or rare (MAF<5%) coding variants that have not been reported previously. Pathway/gene set enrichment analyses of all associated variants highlight lipid particle, adiponectin level, abnormal white adipose tissue physiology, and bone development and morphology as processes affecting fat distribution and body shape. Furthermore, the cross-trait associations and the analyses of variant and gene function highlight a strong connection to lipids, cardiovascular traits, and type 2 diabetes. In functional follow-up analyses, specifically in *Drosophila* RNAi-knockdown crosses, we observed a significant increase in the total body triglyceride levels for two genes (*DNAH10* and *PLXND1*). By examining variants often poorly tagged or entirely missed by genome-wide association studies, we implicate novel genes in fat distribution, stressing the importance of interrogating low-frequency and protein-coding variants.

---

Body fat distribution, as assessed by waist-to-hip ratio (WHR), is a heritable trait and a well-established risk factor for adverse metabolic outcomes^1–6^. A high WHR often indicates a large presence of intra-abdominal fat whereas a low WHR is correlated with a greater accumulation of gluteofemoral fat. Lower values of WHR have been consistently associated with lower risk of cardiometabolic diseases like type 2 diabetes (T2D)^7,8^, or differences in bone structure and gluteal muscle mass^9^. These epidemiological associations are consistent with the results of our previously reported genome-wide association study (GWAS) of 49 loci associated with WHR (after adjusting for body mass index, WHRadjBMI)^10^. Notably, a genetic predisposition to higher WHRadjBMI is associated with increased risk of T2D and coronary heart disease (CHD), and this association appears to be causal^9^.

More recently, large-scale genetic studies have identified ~125 common loci for central obesity, primarily non-coding variants of relatively modest effect, for different measures of body fat distribution^10–16^. Large scale interrogation of both common (minor allele frequency [MAF]≥5%) and low frequency or rare (MAF<5%) coding and splice site variation may lead to additional insights into the genetic and biological etiology of central obesity by narrowing in on causal genes contributing to trait variance. Thus, we set out to identify protein-coding and splice site variants associated with WHRadjBMI using exome array data and to explore their contribution to variation in WHRadjBMI through multiple follow-up analyses.

## RESULTS

### Protein-coding and splice site variation associated with body fat distribution

We conducted a 2-stage fixed-effects meta-analysis testing both additive and recessive models in order to detect protein-coding genetic variants that influence WHRadjBMI (**Online Methods**, Figure 1). Our stage 1 meta-analysis included up to 228,985 variants (218,195 with MAF<5%) in up to 344,369 individuals from 74 studies of European (N=288,492), South Asian (N=29,315), African (N=15,687), East Asian (N=6,800) and Hispanic/Latino (N=4,075) descent, genotyped with an ExomeChip array **(Supplementary Tables 1-3)**. For stage 2, we assessed 70 suggestively significant (*P*<2×l0^−6^) variants from stage 1 in two independent cohorts from the United Kingdom [UK Biobank (UKBB), N=119,572] and Iceland (deCODE, N=12,605) **(Online Methods, Supplementary Data 1-3)** for a total stage 1+2 sample size of 476,546 (88% European). Variants were considered statistically significant in the total meta-analyzed sample (stage 1+2) when they achieved a significance threshold of P<2×l0^−7^ after Bonferroni correction for multiple testing (0.05/246,328 variants tested). Of the 70 variants brought forward, two common and five rare variants were not available in either Stage 2 study **(Tables 1-2, Supplementary Data 1-3)**. Thus, we require P<2×10^−7^ in Stage 1 for significance. Variants are considered novel if they were greater than one megabase (Mb) from a previously-identified WHRadjBMI lead SNP^10–16^.

**Figure 1.**
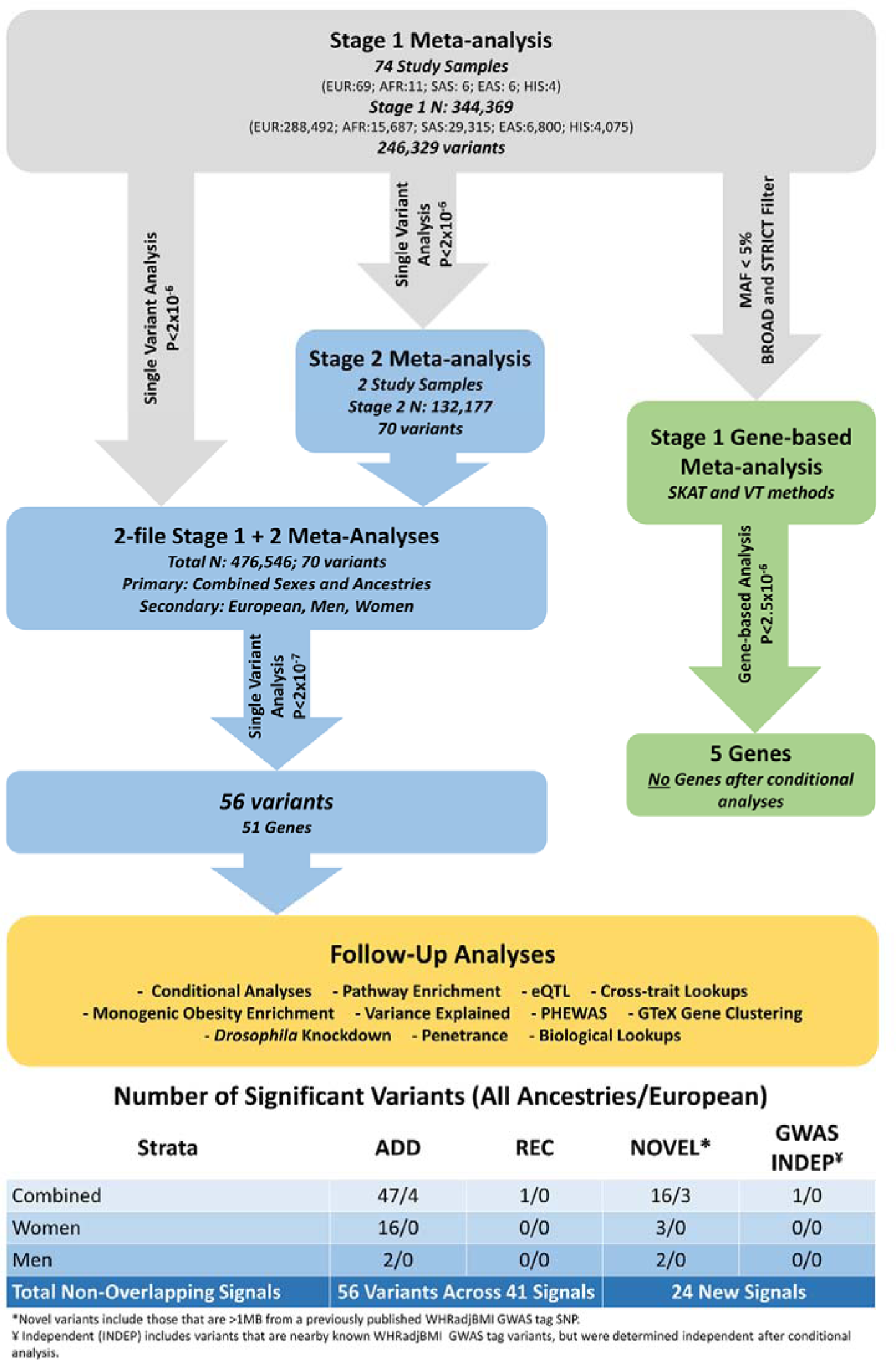
Summary of meta-analysis study design and workflow. Abbreviations: EUR-European, AFR-African, SAS-South Asian, EAS-East Asian, and HIS-Hispanic/Latino ancestry.

In stages 1 and 2 combined all ancestry meta-analyses, we identified 48 coding variants (16 novel) across 43 genes, 47 identified assuming an additive model, and one more variant under a recessive model **(Table 1, Supplementary Figures 1-4)**. Due to the possible heterogeneity introduced by combining multiple ancestries^17^, we also performed a European-only meta-analysis. Here, four additional coding variants were significant (three novel) assuming an additive model **(Table 1, Supplementary Figures 5-8)**. Of these 52 significant variants (48 from the all ancestry and 4 from the European-only analyses), eleven were of low frequency, including seven novel variants in *RAPGEF3, FGFR2, R3HDML, HIST1H1T, PCNXL3, ACVR1C*, and *DARS2*. These low frequency variants tended to display larger effect estimates than any of the previously reported common variants (Figure 2)^10^. In general, variants with MAF<1% had effect sizes approximately three times greater than those of common variants (MAF>5%). Although, we cannot rule out the possibility that additional rare variants with smaller effects sizes exist that, despite our ample sample size, we are still underpowered to detect (See estimated 80% power in Figure 2). However, in the absence of common variants with similarly large effects, our results point to the importance of investigating rare and low frequency variants to identify variants with large effects (Figure 2).

**Figure 2.**
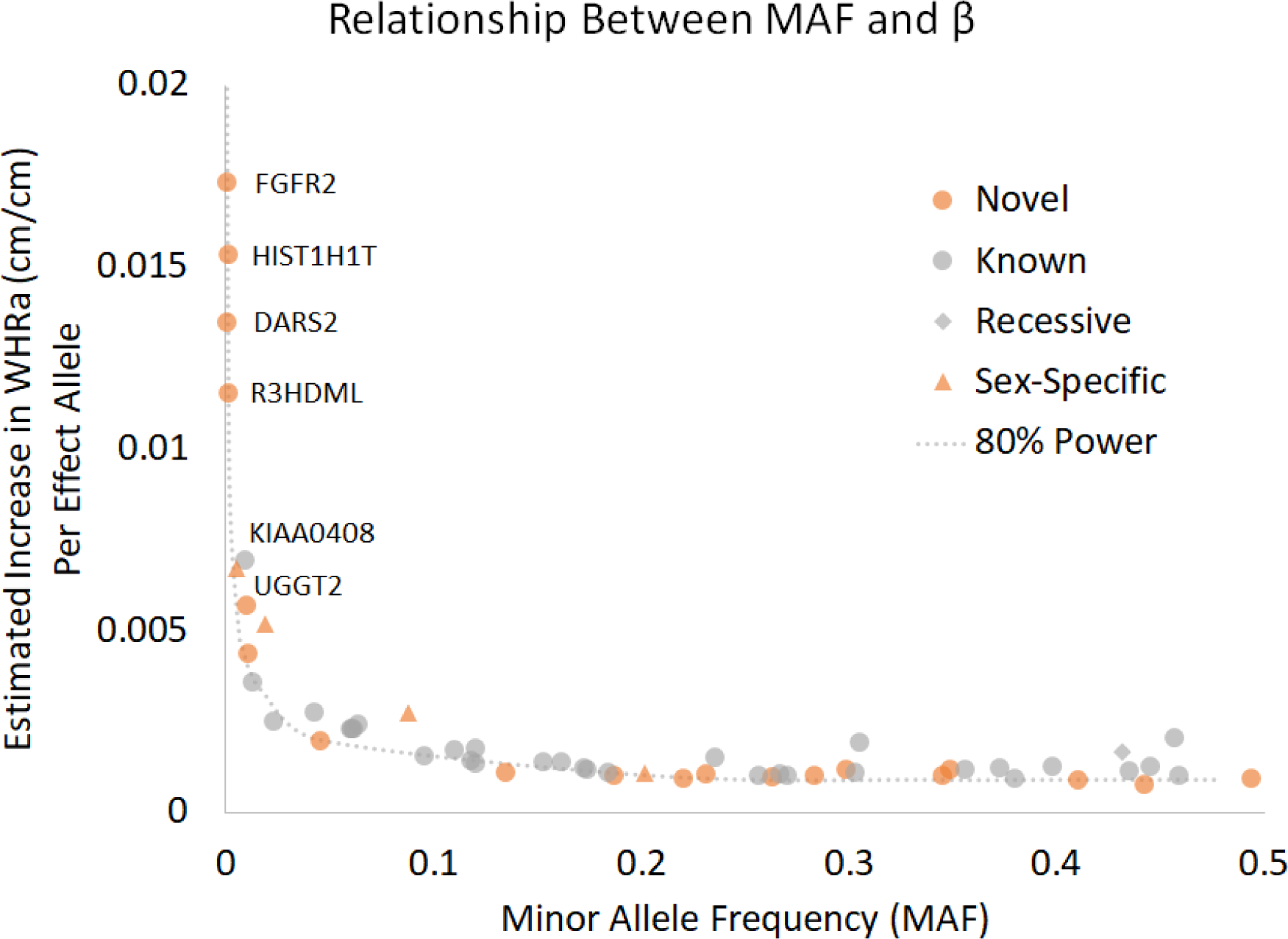
Minor allele frequency compared to estimated effect. This scatter plot displays the relationship between minor allele frequency (MAF) and the estimated effect (β) for each significant coding variant in our meta-analyses. All novel WHRadjBMI variants are highlighted in orange, and variants identified only in models that assume recessive inheritance are denoted by diamonds and only in sex-specific analyses by triangles. Eighty percent power was calculated based on the total sample size in the Stage 1+2 meta-analysis and P-2×10^−7^. Estimated effects are shown in original units (cm/cm) calculated by using effect sizes in standard deviation (SD) units times SD of WHR in the ARIC study (sexes combined=0.067, men=0.052, women=0.080).

Given the established differences in the genetic underpinnings between sexes for WHRadjBMI^10,11^, we also performed sex-stratified analyses and report variants that were array-wide significant (P<2×10^−7^) in at least one sex stratum and exhibit significant sex-specific effects (P_sexhet_<7.14–10^−4^, see **Online Methods**). We found four additional novel variants that were not identified in the sex-combined meta-analyses (in *UGGT2* and *MMP14* for men only; and *DSTYK* and *ANGPTL4* for women only) **(Table 2, Supplementary Figures 9-15)**. Variants in *UGGT2* and *ANGPTL4* were of low frequency (MAF_men_=0.6% and MAF_women_=1.9%, respectively). Additionally, 14 variants from the sex-combined meta-analyses displayed stronger effects in women, including the novel, low frequency variant in *ACVR1C* (rs55920843, MAF=1.1%, **Supplementary Figure 4**). Overall, 19 of the 56 variants (32%) identified across all meta-analyses (48 from all ancestry, 4 from European-only and 4 from sex-stratified analyses) showed significant sex-specific effects on WHRadjBMI (Figure 1): 16 variants with significantly stronger effects in women, and three in men (Figure 1).

In summary, we identified 56 array-wide significant coding variants (P<2.0×10^−7^); 43 common (14 novel) and 13 low frequency or rare variants (9 novel). For all 55 significant variants from the additive model (47 from all ancestry, 4 from European-only, and 4 from sex-specific analyses), we examined potential collider bias^18,19^, i.e. potential bias in effect estimates caused by adjusting for a correlated and heritable covariate like BMI, for the relevant sex stratum and ancestry. We corrected each of the variant - WHRadjBMI associations for the correlation between WHR and BMI and the correlation between the variant and BMI **(Online Methods, Supplementary Table 7, Supplementary Note 1)**. Overall, 51 of the 55 additive model variants were robust against collider bias^18,19^ across all primary and secondary meta-analyses. Of the 55, 25 of the WHRadjBMI variants from the additive model were nominally associated with BMI (P_BMI_<0.05), yet effect sizes changed little after correction for potential biases (15% change in effect estimate on average). For 4 of the 55 SNPs (rsl41845046, rsl034405, rs3617, rs9469913, Table 1), the association with WHRadjBMI appears to be attenuated following correction (P_corrected_> 9–10^−4^, 0.05/55), including one novel variant, rsl034405 in *C30rfl8*. Thus, these 4 variants warrant further functional investigations to quantify their impact on WHR, as a true association may still exist, although the effect may be slightly overestimated in the current analysis.

**Table 1.**
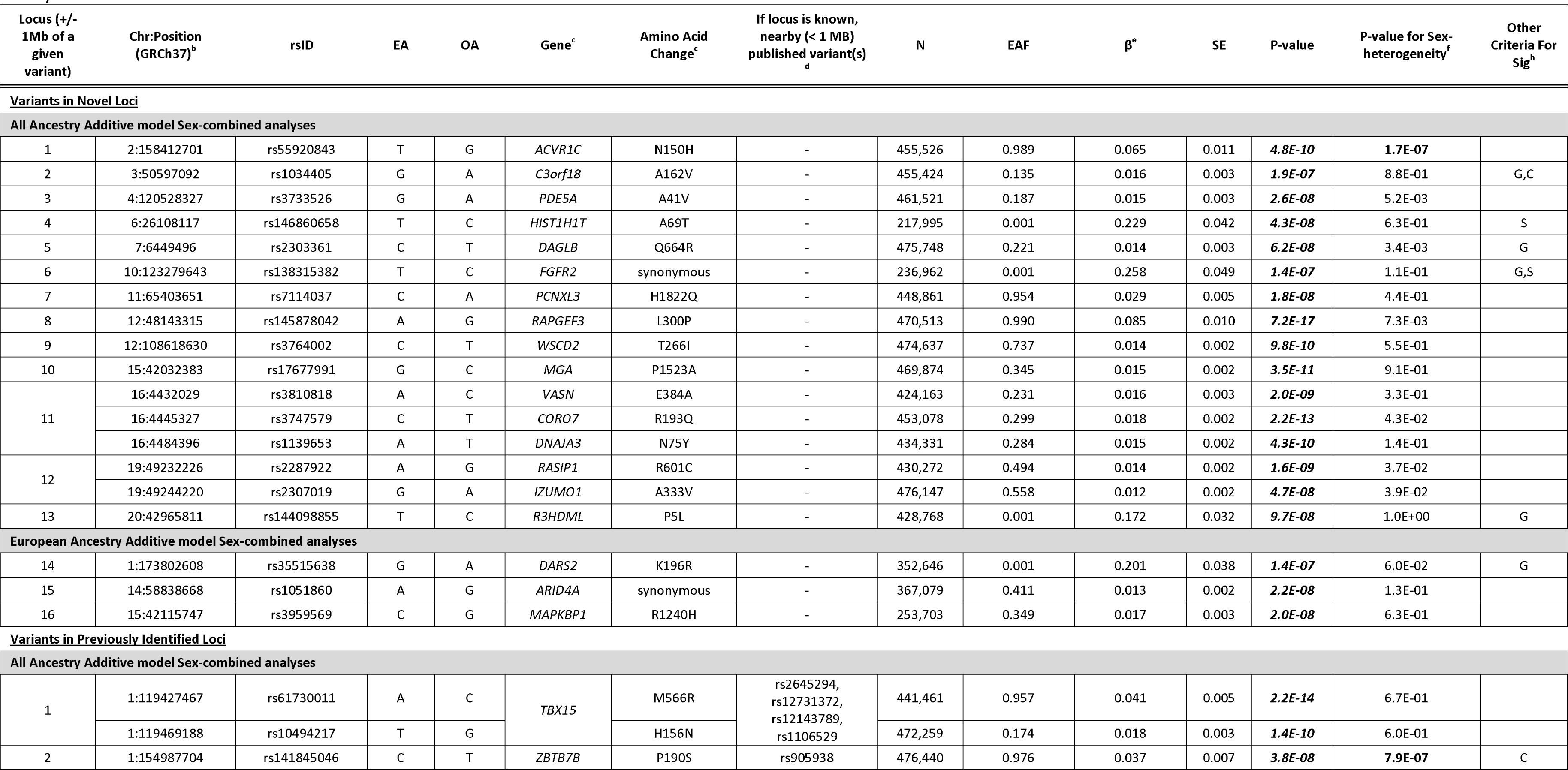
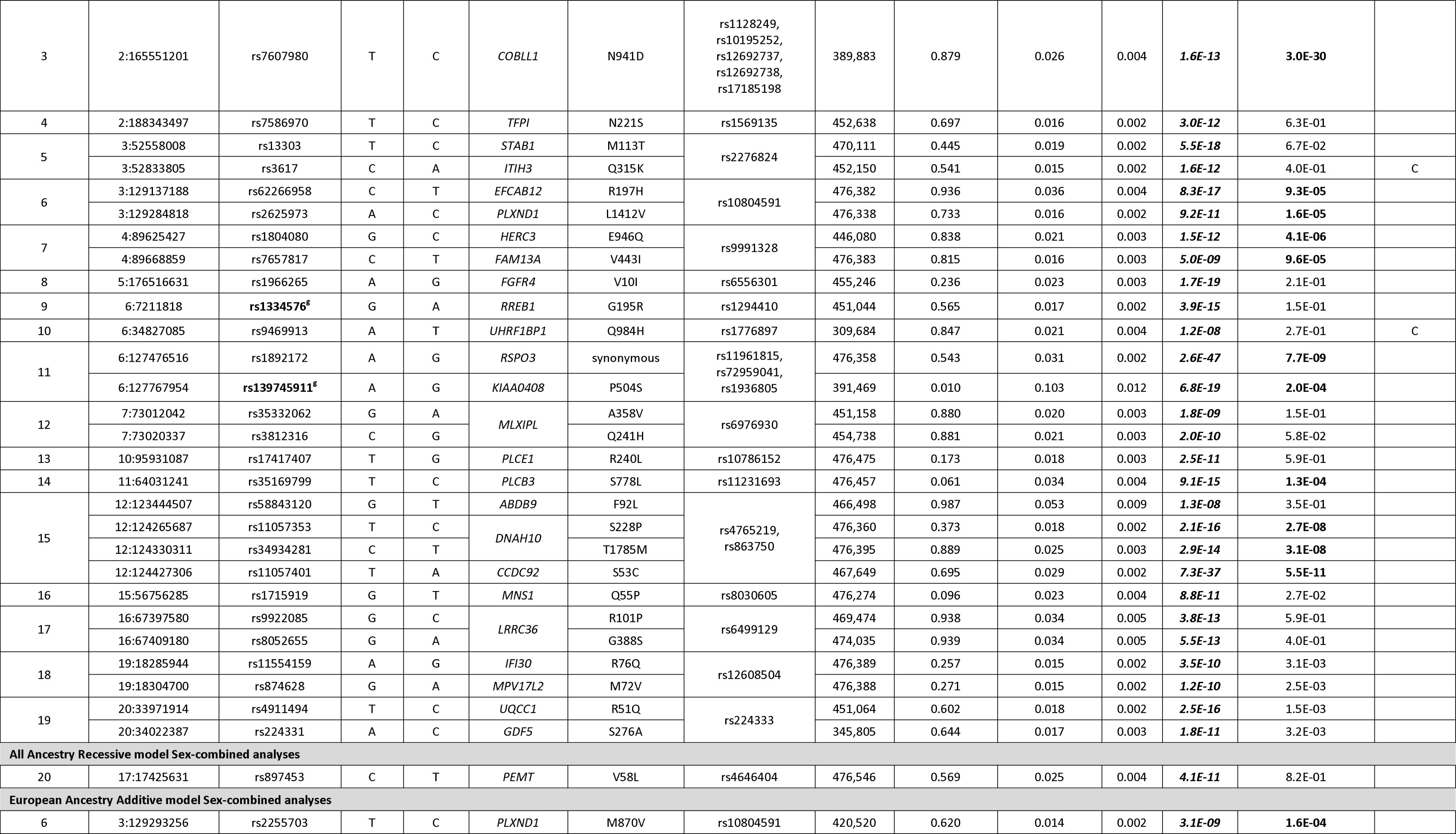
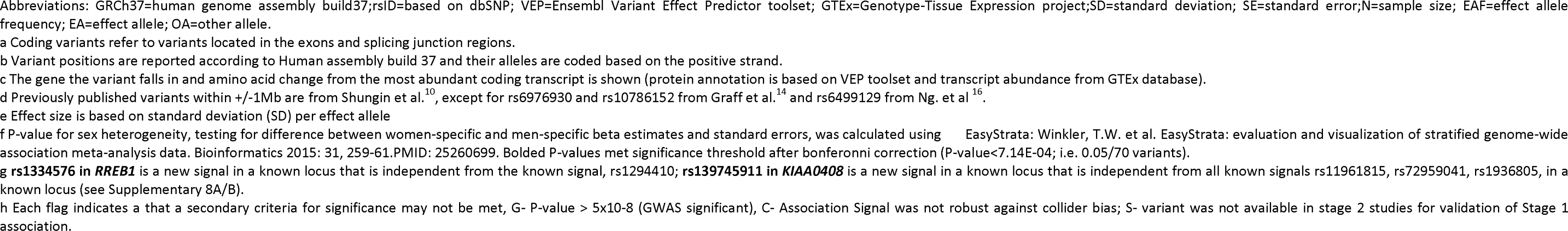
Association results for Combined Sexes. Association results based on an additive or recessive model for coding variants that met array-wide significance (P<2×10-07) in the sex-combined meta-analyses.

Using stage 1 meta-analysis results, we then aggregated low frequency variants across genes and tested their joint effect with both SKAT and burden tests^20^ **(Supplementary Table 8, Online Methods)**. We identified five genes that reached array-wide significance (P<2.5–10^−6^, 0.05/16,222, genes tested), *RAPGEF3, ACVR1C, ANGPTL4, DNAI1*, and *NOP2*. However, while all genes analyzed included more than one variant, none remained significant after conditioning on the single variant with the most significant p-value. We identified variants within *RAPGEF3, ACVR1C, ANGPTL4* that reached suggestive significance in Stage 1 and chip-wide significance in stage 1+2 for one or more meta-analyses (Tables 1 and 2); however, we did not identify any significant variants for *DNAI1* and *NOP2*. While neither of these genes had a single variant that reached chip-wide significance, they each had variants with nearly significant results (NOP2: P=3.69×10^−5^, *DNAI1*: 4.64×10^−5^). Combined effects with these single variants and others in LD within the gene likely drove the association in our aggregate gene-based tests, but resulted in non-significance following conditioning on the top variant. While our results suggest these associations are driven by a single variant, each gene may warrant consideration in future investigations.

**Table 2.**
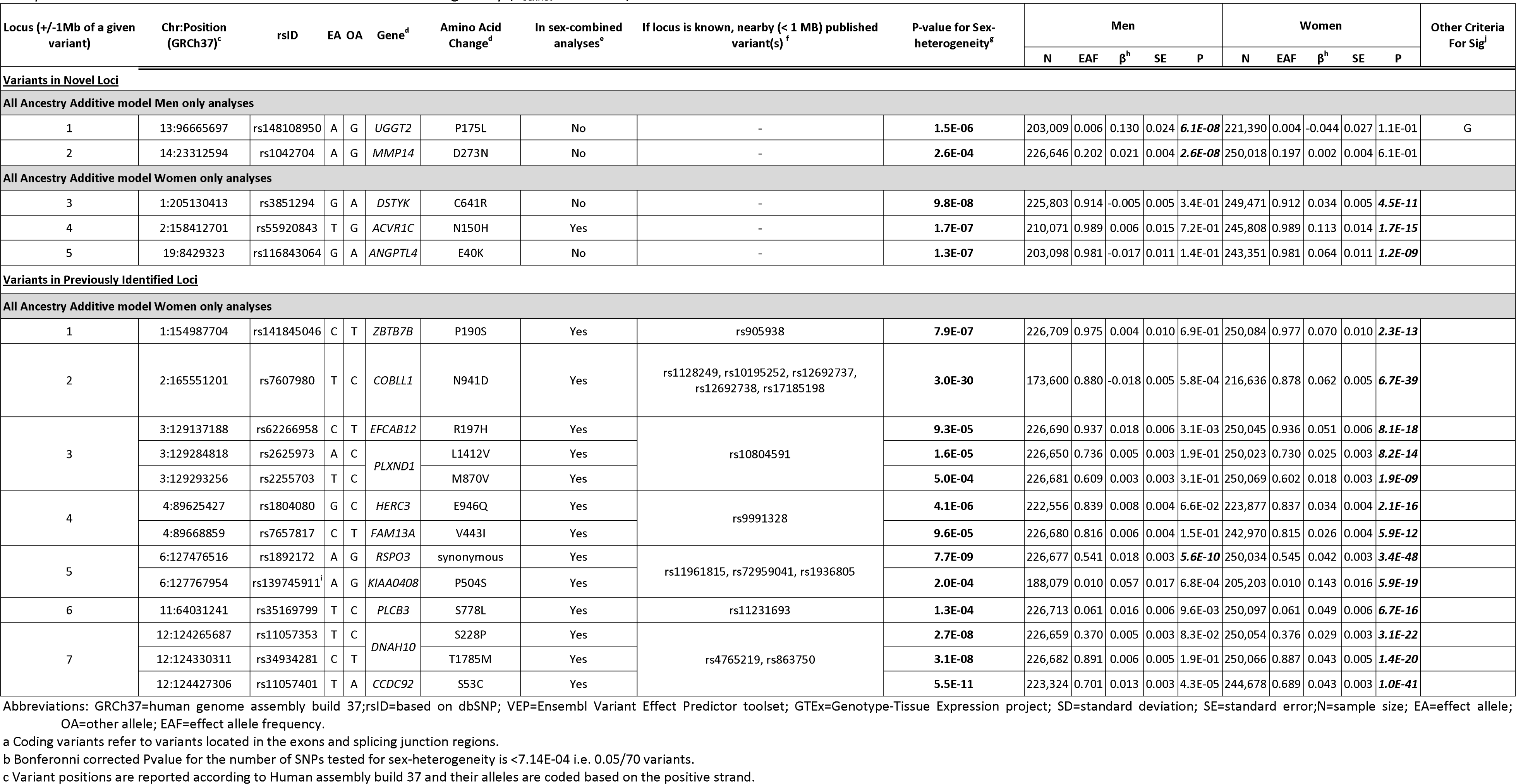
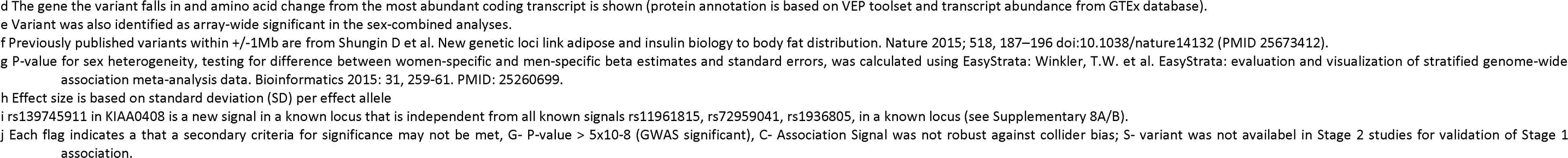
Association results for Sex-stratified analyses. Association results based on an additive or recessive model for coding variants that met array-wide significance (P<2×10-07) in the sex-specific meta-analyses and reach bonferonni corrected P-value for sex hetergeneity (P_seX_het<7.14E-04).

### Conditional analyses

We next implemented conditional analyses to determine (1) the number of independent association signals the 56 array-wide significant coding variants represent, and (2) whether the 33 variants near known GWAS association signals (<+/− 1Mb) represent independent novel association signals. To determine if these variants were independent association signals, we used approximate joint conditional analyses to test for independence in stage 1 **(Online Methods; Supplementary Table 4)**^20^. Only the *RSPO3-KIAA0408* locus contains two independent variants 291 Kb apart, rs1892172 in *RSPO3* (MAF=46.1%, P_conditiona1_=4.37×10^−23^ in the combined sexes, and P_conditionar_=2.4×10^−20^ in women) and rs139745911 in *KIAA0408* (MAF=0.9%, P_conditionar_=3.68×10^−11^ in the combined sexes, and P_conditionarl_=1.46×10^−11^ in women; Figure 3A).

**Figure 3.**
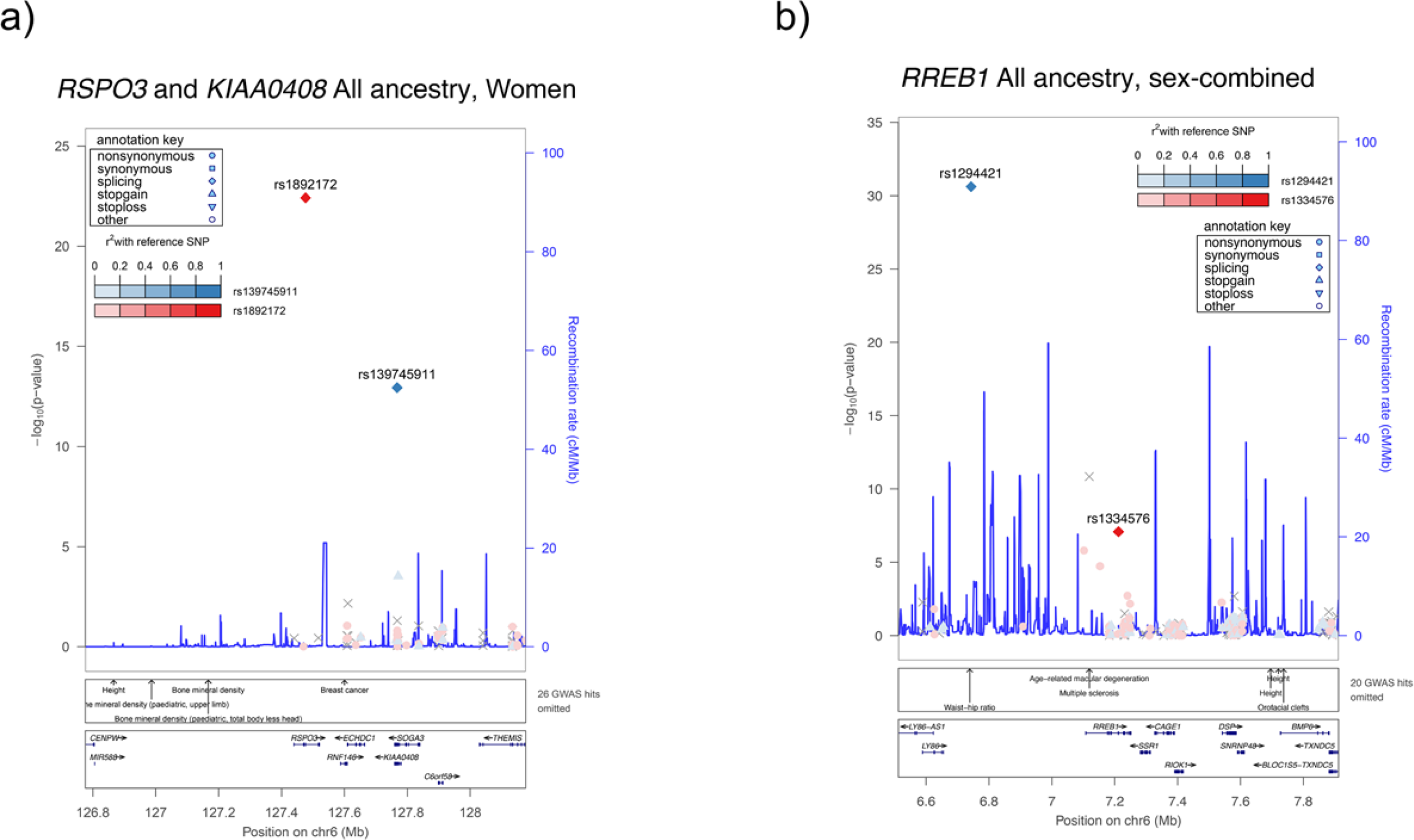
Regional association plots for known loci with novel coding signals. Point color reflects r^2^ calculated from the ARIC dataset. In a) there are two independent variants in *RSP03* and *KIAA0408*, as shown by conditional analysis. In b) we have a variant in *RREB1* that is independent of the GWAS variant rsl294421.

Further, 33 of our significant variants are within one Mb of previously identified GWAS tag SNPs for WHRadjBMI. We again used approximate joint conditional analysis to test for independence in the stage 1 meta-analysis dataset and obtained further complementary evidence from the UKBB dataset where necessary **(Online Methods)**. We identified one coding variant representing a novel independent signal in a known locus [*RREB1*; stage1 meta-analysis, rs1334576, EAF = 0.44, P_conditionar_=3.06×10^−7^, **(Supplementary Table 5, Figure 3 [B])**; UKBB analysis, rs1334576, *RREB1*, P_conditionar_=1.24×l0^−8^, **(Supplementary Table 6)** in the sex-combined analysis.

In summary, we identified a total of 56 WHRadjBMI-associated coding variants in 41 independent association signals. Of these 41 independent association signals, 24 are new or independent of known GWAS-identified tag SNPs (either >1MB +/− or array-wide significant following conditional analyses) (Figure 1). Thus, bringing our total to 15 common and 9 low-frequency or rare novel variants following conditional analyses. The remaining non-GWAS-independent variants may assist in narrowing in on the causal variant or gene underlying these established association signals.

### Gene set and pathway enrichment analysis

To determine if the significant coding variants highlight novel biological pathways and/or provide additional support for previously identified biological pathways, we applied two complementary pathway analysis methods using the EC-DEPICT (ExomeChip Data-driven Expression Prioritized Integration for Complex Traits) pathway analysis tool,^21,22^ and PASCAL^23^ **(Online Methods)**. While for PASCAL all variants were used, in the case of EC-DEPICT, we examined 361 variants with suggestive significance (P<5×10^−4^)^10,17^ from the combined ancestries and combined sexes analysis (which after clumping and filtering became 101 lead variants in 101 genes). We separately analyzed variants that exhibited significant sex-specific effects (P_sexhet_<5×10^−4^).

The sex-combined analyses identified 49 significantly enriched gene sets (FDR<0.05) that grouped into 25 meta-gene sets **(Supplementary Note 2, Supplementary Data 4-5)**. We noted a cluster of meta-gene sets with direct relevance to metabolic aspects of obesity (“enhanced lipolysis,” “abnormal glucose homeostasis,” “increased circulating insulin level,” and “decreased susceptibility to diet-induced obesity”); we observed two significant adiponectin-related gene sets within these meta-gene sets. While these pathway groups had previously been identified in the GWAS DEPICT analysis (Figure 4), many of the individual gene sets within these meta-gene sets were not significant in the previous GWAS analysis, such as “insulin resistance,” “abnormal white adipose tissue physiology,” and “abnormal fat cell morphology” **(Supplementary Data 4, Figure 4, Supplementary Figure 16a)**, but represent similar biological underpinnings implied by the shared meta-gene sets. Despite their overlap with the GWAS results, these analyses highlight novel genes that fall outside known GWAS loci, based on their strong contribution to the significantly enriched gene sets related to adipocyte and insulin biology (e.g. *MLXIPL, ACVR1C, and ITIH5)* (Figure 4).

**Figure 4.**
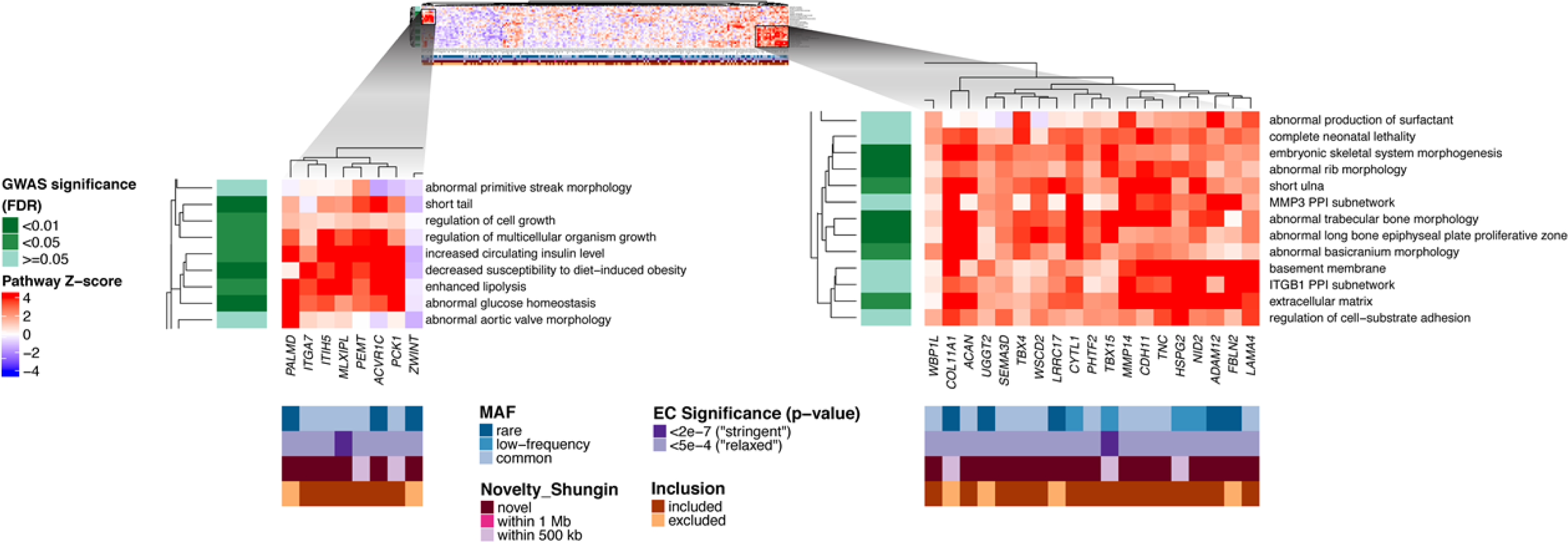
Heat maps showing DEPICT gene set enrichment results. For any given square, the color indicates how strongly the corresponding gene (shown on the x-axis) is predicted to belong to the reconstituted gene set (y-axis). This value is based on the gene’s z-score for gene set inclusion in DEPICT’s reconstituted gene sets, where red indicates a higher and blue a lower z-score. To visually reduce redundancy and increase clarity, we chose one representative “meta-gene set” for each group of highly correlated gene sets based on affinity propagation clustering **(Online Methods, Supplementary Note 2)**. Heatmap intensity and DEPICT P-values (see P-values in **Supplementary Data 4-5)** correspond to the most significantly enriched gene set within the meta-gene set. Annotations for the genes indicate (1) the minor allele frequency of the significant ExomeChip (EC) variant (shades of blue; if multiple variants, the lowest-frequency variant was kept), (2) whether the variant’s P-value reached array-wide significance (<2×10^−7^) or suggestive significance (<5×10^−4^) (shades of purple), (3) whether the variant was novel, overlapping “relaxed” GWAS signals from Shungin et al.^10^ (GWAS P<5xl0^−4^), or overlapping “stringent” GWAS signals (GWAS P<5xl0^−8^) (shades of pink), and (4) whether the gene was included in the gene set enrichment analysis or excluded by filters (shades of brown/orange) (Online Methods and Supplementary Information). Annotations for the gene sets indicate if the meta-gene set was found significant (shades of green; FDR <0.01, <0.05, or not significant) in the DEPICT analysis of GWAS results from Shungin et al.

To focus on novel findings, we conducted pathway analyses after excluding variants from previous WHRadjBMI analyses^10^ **(Supplemental Note 2)**. Seventy-five loci/genes were included in the EC-DEPICT analysis, and we identified 26 significantly enriched gene sets (13 meta-gene sets). Here, all but one gene set, “lipid particle size”, were related to skeletal biology. This result likely reflects an effect on the pelvic skeleton (hip circumference), shared signaling pathways between bone and fat (such as TGF-beta) and shared developmental origin^24^ **(Supplementary Data 5, Supplementary Figure 16b)**. Many of these pathways were previously found to be significant in the GWAS DEPICT analysis; these findings provide a fully independent replication of their biological relevance for WHRadjBMI.

We used PASCAL **(Online Methods)** to further distinguish between enrichment based on *coding-only* variant associations (this study) and *regulatory-only* variant associations (up to 20 kb upstream of the gene from a previous GIANT study^10^). For completeness, we also compared the coding pathways to those that could be identified in the total previous GWAS effort (using both *coding and regulatory* variants) by PASCAL. The analysis revealed 116 significantly enriched coding pathways (FDR<0.05; **Supplementary Table 9**). In contrast, a total of 158 gene sets were identified in the coding+regulatory analysis that included data from the previous GIANT waist GWAS study. Forty-two gene sets were enriched in both analyses. Thus, while we observed high concordance in the −log10 (p-values) between ExomeChip and GWAS gene set enrichment (Pearson’s r (coding *vs* regulatory only) = 0.38, P<10^−300^; Pearson’s r (coding *vs* coding+regulatory)=0.51, P<10^−300^), there are gene sets that seem to be enriched *specifically* for variants in coding regions (e.g., decreased susceptibility to diet-induced obesity, abnormal skeletal morphology) or unique to variants in regulatory regions (e.g. transcriptional regulation of white adipocytes) **(Supplementary Figure 17)**.

The EC-DEPICT and PASCAL results showed a moderate but strongly significant correlation (for EC-DEPICT and the PASCAL max statistic, r=.277 with ρ=9.8×10^−253^; for EC-DEPICT and the PASCAL sum statistic, r=.287 with p = 5.42×10^−272^). Gene sets highlighted by both methods strongly implicated a role for pathways involved in skeletal biology, glucose homeostasis/insulin signaling, and adipocyte biology. Indeed, we are even more confident in the importance of this core overlapping group of pathways due to their discovery by both methods **(Supplementary Figure 18)**.

### Cross-trait associations

To assess the relevance of our identified variants with cardiometabolic, anthropometric, and reproductive traits, we conducted association lookups from existing ExomeChip studies of 15 traits **(Supplementary Data 6, Supplementary Figure 19)**. Indeed, the clinical relevance of central adiposity is likely to be found in the cascade of impacts such variants have on downstream cardiometabolic disease.^22,25–29^ We found that variants in *STAB1* and *PLCB3* display the greatest number of significant cross-trait associations, each associating with seven different traits (P<9.8×10^−4^, 0.05/51 variants tested). Of note, these two genes cluster together with *RSPO3, DNAH10, MNS1, COBLL1, CCDC92*, and *ITIH3* **(Supplementary Data 6, Supplementary Figure 19)**. The WHR-increasing alleles in this cluster of variants exhibit a pattern of increased cardiometabolic risk (e.g. increased fasting insulin [FI], two-hour glucose [TwoHGlu], and triglycerides [TG]; and decreased high-density lipoprotein cholesterol [HDL]), but also decreased BMI. This phenomenon, where variants associated with lower BMI are also associated with increased cardiometabolic risk, has been previously reported.^30–36^. A recent Mendelian Randomization (MR) analysis of the relationship between central adiposity (measured as WHRadjBMI) and cardiometabolic risk factors found central adiposity to be causal.^9^ Using 48 WHR-increasing variants reported in the recent GIANT analysis^10^ to calculate a polygenic risk score, Emdin *et al.* found that a 1 SD increase in genetic risk of central adiposity was associated with higher total cholesterol, triglyceride levels, fasting insulin and two-hour glucose, and lower HDL - all indicators of cardiometabolic disease, and also associated with a 1 unit decrease in BMI^9^.

We conducted a search in the NHGRI-EBI GWAS Catalog^37,38^ to determine if any of our significant ExomeChip variants are in high LD (R^2^>0.7) with variants associated with traits or diseases not covered by our cross trait lookups **(Supplementary Data 7)**. We identified several cardiometabolic traits (adiponectin, coronary heart disease *etc.*) and behavioral traits potentially related to obesity (carbohydrate, fat intake *etc.*) with GWAS associations that were not among those included in cross-trait analyses and nearby one or more of our WHRadjBMI-associated coding variants. Additionally, many of our ExomeChip variants are in LD with GWAS variants associated with other behavioral and neurological traits (schizophrenia, bipolar disorder etc.), and inflammatory or autoimmune diseases (Crohn’s Disease, multiple sclerosis *etc.*) **(Supplementary Data 7)**.

Given the established correlation between total body fat percentage and WHR (R=0.052 to 0.483)^39–41^, we examined the association of our top exome variants with both total body fat percentage (BF%) and truncal fat percentage (TF%) available in a sub-sample of up to 118,160 participants of UKBB **(Supplementary Tables 10-11)**. Seven of the common novel variants were significantly associated (P<0.001, 0.05/48 variants examined) with both BF% and TF% in the sexes-combined analysis (*COBLL1, UHRF1BP1, WSCD2, CCDC92, IFI30, MPV17L2, IZUMO1)*. Only one of our tag SNPs, rs7607980 in *COBLL1*, is nearby a known total body for percentageBF% GWAS locus (rs6738627; *R*^*2*^=0.1989, distance=6751 bp, with our tag SNP)^42^. Two additional variants, rs62266958 in *EFCAB12* and rs224331 in *GDF5*, were significantly associated with TF% in the women-only analysis. Of the nine SNPs associated with at least one of these two traits, all variants displayed much greater magnitude of effect on TF% compared to BF% **(Supplementary Figure 20)**.

Previous studies have demonstrated the importance of examining common and rare variants within genes with mutations known to cause monogenic diseases^43,44^. We assessed enrichment of our WHRadjBMI within genes that cause monogenic forms of lipodystrophy) and/or insulin resistance **(Supplementary Data 8)**. No significant enrichment was observed **(Supplementary Figure 21)**. For lipodystrophy, the lack of significant findings may be due in part to the small number of implicated genes and the relatively small number of variants in monogenic disease-causing genes, reflecting their intolerance of variation.

### Genetic architecture of WHRadjBMI coding variants

We used summary statistics from our stage 1 results to estimate the phenotypic variance explained by ExomeChip coding variants. We calculated the variance explained by subsets of SNPs across various significance thresholds (P< 2×10^−7^ to 0.2) and conservatively estimated using only independent tag SNPs **(Supplementary Table 12, Online Methods,** and **Supplementary Figure 22)**. The 22 independent significant coding SNPs in stage 1 account for 0.28% of phenotypic variance in WHRadjBMI. For independent variants that reached suggestive significance in stage 1 (P<2×10^−6^), 33 SNPs explain 0.38% of the variation; however, the 1,786 independent SNPs with a liberal threshold of P<0.02 explain 13 times more variation (5.12%). While these large effect estimates may be subject to winner’s curse, for array-wide significant variants, we detected a consistent relationship between effect magnitude and MAF in our stage 2 analyses in UK Biobank and deCODE **(Supplementary Data 1-3)**. Notably, the Exomechip coding variants explained less of the phenotypic variance than in our previous GIANT investigation, wherein 49 significant SNPs explained 1.4% of the variance in WHRadjBMI. When considering all coding variants on the ExomeChip in men and women together, 46 SNPs with a P<2×10^−6^ and 5,917 SNPs with a P<0.02 explain 0.51% and 13.75% of the variance in WHRadjBMI, respectively. As expected given the design of the ExomeChip, the majority of the variance explained is attributable to rare and low frequency coding variants (independent SNPs with MAF<1% and MAF<5% explain 5.18% and 5.58%, respectively). However, for rare and low frequency variants, those that passed significance in stage 1 explain only 0.10% of the variance in WHRadjBMI. As in Figure 2, these results also indicate that there are additional coding variants associated with WHRadjBMI that remain to be discovered, particularly rare and low frequency variants with larger effects than common variants. Due to observed differences in association strength between women and men, we estimated variance explained for the same set of SNPs in women and men separately. As observed in previous studies^10^, there was significantly (P_RsqDiff_<0.002=0.05/21, Bonferroni-corrected threshold) more variance explained in women compared to men at each significance threshold considered (differences ranged from 0.24% to 0.91%).

To better understand the potential clinical impact of WHRadjBMI associated variants, we conducted penetrance analysis using the UKBB population (both sexes combined, and men- and women-only). We compared the number of carriers and non-carriers of the minor allele for each of our significant variants in centrally obese and non-obese individuals to determine if there is a significant accumulation of the minor allele in either the centrally obese or non-obese groups **(Online Methods)**. Three rare and low frequency variants (MAF ≤ 1%) with larger effect sizes (effect size > 0.90) were included in the penetrance analysis using World Health Organization (WHO-obese women WHR>0.85 and obese men WHR>0.90) WHR cut-offs for central obesity. Of these, one SNV (rs55920843-ACVR1C; P_sex-combined_=9.25×10^−5^; P_women_=4.85×10^−5^) showed a statistically significant difference in the number of carriers and non-carriers of the minor allele when the two strata were compared (sex-combined obese carriers=2.2%; non-obese carriers=2.6%; women obese carriers=2.1%; non-obese women carriers=2.6% **(Supplementary Table 13, Supplementary Figure 23)**. These differences were significant in women, but not in men (P_men_<5.5×10^−3^ after Bonferroni correction for 9 tests) and agree with our overall meta-analysis results, where the minor allele (G) was significantly associated with lower WHRadjBMI in women only (Tables 1 and 2).

### Evidence for functional role of significant variants

#### Drosophila Knockdown

Considering the genetic evidence of adipose and insulin biology in determining body fat distribution^10^, and the lipid signature of the variants described here, we examined whole-body triglycerides levels in adult *Drosophila*, a model organism in which the fat body is an organ functionally analogous to mammalian liver and adipose tissue and triglycerides are the major source of fat storage^45^. Of the 51 genes harboring our 56 significantly associated variants, we identified 27 with *Drosophila* orthologues for functional follow-up analyses. In order to prioritize genes for follow-up, we selected genes with large changes in triglyceride storage levels (> 20% increase or > 40% decrease, as chance alone is unlikely to cause changes of this magnitude, although some decrease is expected) after considering each corresponding orthologue in an existing large-scale screen for adipose with ≤2 replicates per knockdown strain.^45^ Two orthologues, for *PLXND1* and *DNAH10*, from two separate loci met these criteria. For these two genes, we conducted additional knockdown experiments with ≥5 replicates using tissue-specific drivers (fat body [cg-Gal4] and neuronal [elav-Gal4] specific RNAi-knockdowns) **(Supplementary Table 14)**. A significant (P<0.025, 0.05/2 orthologues) increase in the total body triglyceride levels was observed in *DNAH10* orthologue knockdown strains for both the fat body and neuronal drivers. However, only the neuronal driver knockdown for *PLXND1* produced a significant change in triglyceride storage. *DNAH10* and *PLXND1* both lie within previous GWAS identified regions. Adjacent genes have been highlighted as likely candidates for the *DNAH10* association region, including *CCDC92* and *ZNF664* based on eQTL evidence. However, our fly knockdown results support *DNAH10* as the causal genes underlying this association. Of note, rsll057353 in *DNAH10* showed suggestive significance after conditioning on the known GWAS variants in nearby *CCDC92* (sex-combined P_conditional_=7.56×10^−7^; women-only rsll057353 P_conditional_=5.86×10^−7^, **Supplementary Table 6**; thus providing some evidence of multiple causal variants/genes underlying this association signal. Further analyses are needed to determine whether the implicated coding variants from the current analysis are the putatively functional variants, specifically how these variants affect transcription in and around these loci, and exactly how those effects alter biology of relevant human metabolic tissues.

#### eQTL Lookups

To gain a better understanding of the potential functionality of novel and low frequency variants, we examined the *cis*-association of the identified variants with expression level of nearby genes in subcutaneous adipose tissue, visceral omental adipose tissue, skeletal muscle and pancreas from GTEx^46^, and assessed whether the exome and eQTL associations implicated the same signal **(Online Methods, Supplementary Data 9, Supplementary Table 15)**. The lead exome variant was associated with expression level of the coding gene itself for *DAGLB, MLXIPL, CCDC92, MAPKBP1, LRRC36* and *UQCC1*. However, at three of these loci (*MLXIPL, MAPKBP1*, and *LRRC36*), the lead exome variant is also associated with expression level of additional nearby genes, and at three additional loci, the lead exome variant is only associated with expression level of nearby genes (*HEMK1* at *C3orf18; NT5DC2, SMIM4* and *TMEM110* at *STAB1/ITIH3;* and *C6orf106* at *UHRF1BP1*). Although detected with a missense variant, these loci are also consistent with a regulatory mechanism of effect as they are significantly associated with expression levels of genes, and the association signal may well be due to LD with nearby regulatory variants.

Some of the coding genes implicated by eQTL analyses are known to be involved in adipocyte differentiation or insulin sensitivity: e. g. for *MLXIPL*, the encoded carbohydrate responsive element binding protein is a transcription factor, regulating glucose-mediated induction of de *novo* lipogenesis in adipose tissue, and expression of its beta-isoform in adipose tissue is positively correlated with adipose insulin sensitivity^47,48^. For *CCDC92*, the reduced adipocyte lipid accumulation upon knockdown confirmed the involvement of its encoded protein in adipose differentiation^49^.

#### Biological Curation

To gain further insight into the possible functional role of the identified variants, we conducted thorough searches of the literature and publicly available bioinformatics databases **(Supplementary Data 10-11, Box 1, Online Methods)**. Many of our novel low frequency variants are in genes that are intolerant of nonsynonymous mutations (e.g. *ACVR1C, DARS2, FGFR2*; ExAC Constraint Scores >0.5). Like previously identified GWAS variants, several of our novel coding variants lie within genes that are involved in glucose homeostasis (e.g. *ACVR1C, UGGT2, ANGPTL4*), angiogenesis (*RASIP1*), adipogenesis (*RAPGEF3*), and lipid biology (*ANGPTL4, DAGLB*) **(Supplementary Data 10, Box 1)**.

## DISCUSSION

Our two-staged approach to analysis of coding variants from ExomeChip data in up to 476,546 individuals identified a total of 56 array-wide significant variants in 41 independent association signals, including 24 newly identified (23 novel and one independent of known GWAS signals) that influence WHRadjBMI. Nine of these variants were low frequency or rare, indicating an important role for low frequency variants in the polygenic architecture of fat distribution and providing further insights into its underlying etiology. While, due to their rarity, these coding variants only explain a small proportion of the trait variance at a population level, they may, given their predicted role, be more functionally tractable than non-coding variants and have a critical impact at the individual and clinical level. For instance, the association between a low frequency variant (rsll209026; R381Q; MAF<5% in ExAC) located in the *IL23R* gene and multiple inflammatory diseases (such as psoriasis^50^, rheumatoid arthritis^51^, ankylosing spondylitis^52^, and inflammatory bowel diseases^53^) led to the development of new therapies, targeting *IL23* and *IL12* in the same pathway (reviewed in ^54–56^). Thus, we are encouraged that our associated low frequency coding variants displayed large effect sizes; all but one of the nine novel low frequency variants had an effect size larger than the 49 SNPs reported in Shungin *et al.* 2015, and some of these effect sizes were up to 7-fold larger than those previously reported for GWAS. This finding mirrors results for other cardiometabolic traits^57^, and suggests variants of possible clinical significance with even larger effect and lower frequency variants will likely be detected through larger additional genome-wide scans of many more individuals.

We continue to observe sexual dimorphism in the genetic architecture of WHRadjBMI^11^. Overall, we identified 19 coding variants that display significant sex differences, of which 16 (84%) display larger effects in women compared to men. Of the variants outside of GWAS loci, we reported three (two with MAF<5%) that show a significantly stronger effect in women and two (one with MAF<5%) that show a stronger effect in men. Additionally, genetic variants continue to explain a higher proportion of the phenotypic variation in body fat distribution in women compared to men^10,11^. Of the novel female (*DSTYK* and *ANGPTL4*) and male (*UGGT2* and *MMP14*) specific signals, only *ANGPTL4* implicated fat distribution related biology associated with both lipid biology and cardiovascular traits (Box 1). Sexual dimorphism in fat distribution is apparent from childhood and throughout adult life^58–60^, and at sexually dimorphic loci, hormones with different levels in men and women may interact with genomic and epigenomic factors to regulate gene activity, though this remains to be experimentally documented. Dissecting the underlying molecular mechanisms of the sexual dimorphism in body fat distribution, and also how it is correlated with - and causing - important comorbidities like T2D and cardiovascular diseases will be crucial for improved understanding of disease risk and pathogenesis.

### Box 1. Genes of biological interest harboring WHR-associated variants

***PLXND1-*** (3:129284818, rs2625973, known locus) The major allele of a common non-synonymous variant in Plexin Dl (L1412 V, MAF=26.7%) is associated with increased WHRadjBMI (β (SE)= 0.0156 (0.0024), P-value=9.16×10^−11^). *PLXND1* is a semaphorin class 3 and 4 receptor gene, and therefore, is involved in cell to cell signaling and regulation of growth in development for a number of different cell and tissue types, including those in the cardiovascular system, skeleton, kidneys, and the central nervous system^97–101^. Mutations in this gene are associated with Moebius syndrome^102–105^, and persistent truncus arteriosus^99,106^. *PLXND1* is involved in angiogenesis as part of the SEMA and VEGF signalling pathways^107–110^. *PLXND1* was implicated in the development of T2D through its interaction with *SEMA3E* in mice. *SEMA3E* and *PLXND1* are upregulated in adipose tissue in response to diet-induced obesity, creating a cascade of adipose inflammation, insulin resistance, and diabetes mellitus^101^. *PLXND1* is highly expressed in adipose (both subcutaneous and visceral) (GTeX). *PLXND1* is highly intolerant of mutations and therefore highly conserved **(Supplementary Data 10).** Last, our lead variant is predicted as damaging or possibly damaging for all algorithms examined (SIFT, Polyphen2/HDIV, Polyphen2/HVAR, LRT, MutationTaster).

***ACVR1C-*** (2:158412701, rs55920843, novel locus) The major allele of a low frequency non-synonymous variant in activin A receptor type IC (rs55920843, N150H, MAF=1.1%) is associated with increased WHRadjBMI (β (SE)= 0.0652 (0.0105), P-value= 4.81×10^−10^). *ACVR1C*, also called Activin receptor-like kinase 7 (*ALK7*), is a type I receptor for TGFB (Transforming Growth Factor, Beta-1), and is integral for the activation of SMAD transcription factors; therefore, *ACVR1C* plays an important role in cellular growth and differentiation^64–68^, including adipocytes^68^. Mouse Acvrlc decreases secretion of insulin and is involved in lipid storage^69,72,73,69,72,73,111^. *ACVR1C* exhibits the highest expression in adipose tissue, but is also highly expressed in the brain (GTEx)^69–71^. Expression is associated with body fat, carbohydrate metabolism and lipids in both obese and lean individuals^70^. *ACVRlC* is moderately tolerant of mutations (EXaC Constraint Scores: synonymous= −0.86, nonsynonymous = 1.25, LoF = 0.04, **Supplementary Data 10**). Last, our lead variant is predicted as damaging for two of five algorithms examined (LRT and MutationTaster).

***FGFR2-*** (10:123279643, rsl38315382, novel locus) The minor allele of a rare synonymous variant in Fibroblast Growth Factor Receptor 2 (rsl38315382, MAF=0.09%) is associated with increased WHRadjBMI (β (SE) = 0.258 (0.049), P-value= 1.38×10^−07^). The extracellular portion of the FGFR2 protein binds with fibroblast growth factors, influencing mitogenesis and differentiation. Mutations in this gene have been associated with many rare monogenic disorders, including skeletal deformities, craniosynostosis, eye abnormalities, and LADD syndrome, as well as several cancers including breast, lung, and gastric cancer. Methylation of *FGFR2* is associated with high birth weight percentile^112^. *FGFR2* is tolerant of synonymous mutations, but highly intolerant of missense and loss-of-function mutations (ExAC Constraint scores: synonymous=−0.9, missense=2.74, LoF=1.0, **Supplementary Data 10**). Last, this variant is not predicted to be damaging based on any of the 5 algorithms tested.

***ANGPTL4-*** (19:8429323, rsll6843064, novel locus) The major allele of a nonsynonymous low frequency variant in Angiopoietin Like 4 (rsll6843064, E40K, EAF=98.1%) is associated with increased WHRadjBMI (β (SE) = 0.064 (0.011) P-value= 1.20×10^−09^). *ANGPTL4* encodes a glycosylated, secreted protein containing a C-terminal fibrinogen domain. The encoded protein is induced by peroxisome proliferation activators and functions as a serum hormone that regulates glucose homeostasis, triglyceride metabolism^113,114^, and insulin sensitivity^115^. Angptl4-deficient mice have hypotriglyceridemia and increased lipoprotein lipase (LPL) activity, while transgenic mice overexpressing Angplt4 in the liver have higher plasma triglyceride levels and decreased LPL activity^116^. The major allele of rsll6843064 has been previously associated with increased risk of coronary heart disease and increased TG^63^. *ANGPTL4* is moderately tolerant of mutations (ExAC constraint scores synonymous=1.18, missense=0.21, LoF=0.0, **Supplementary Data 10**). Last, our lead variant is predicted damaging for four of five algorithms (SIFT, Polyphen 2/HDIV, Polyphen2/HVAR, and MutationTaster).

***RREB1-*** (6:7211818, rsl334576, novel association signal) The major allele of a common non-synonymous variant in the Ras responsive element binding protein 1 (rsl334576, G195R, EAF=56%) is associated with increased WHRadjBMI (β (SE)=0.017 (0.002), P-value=3.9×l0^−15^). This variant is independent of the previously reported GWAS signal in the *RREB1* region (rsl294410; 6:6738752^10^). The protein encoded by this gene is a zinc finger transcription factor that binds to RAS-responsive elements (RREs) of gene promoters. It has been shown that the calcitonin gene promoter contains an RRE and that the encoded protein binds there and increases expression of calcitonin, which may be involved in Ras/Raf-mediated cell differentiation^117–119^. The ras responsive transcription factor *RREB1* is a candidate gene for type 2 diabetes associated end-stage kidney disease^118^. This variant is highly intolerant to loss of function (ExAC constraint score LoF = 1, **Supplementary Data 10**).

***DAGLB-*** (7:6449496, rs2303361, novel locus) The minor allele of a common non-synonymous variant (rs2303361, Q664R, MAF=22%) in *DAGLB* (Diacylglycerol lipase beta) is associated with increased WHRadjBMI (β (SE)= 0.0136 (0.0025), P-value=6.24×10^−8^). *DAGLB* is a diacylglycerol (DAG) lipase that catalyzes the hydrolysis of DAG to 2-arachidonoyl-glycerol, the most abundant endocannabinoid in tissues. In the brain, DAGL activity is required for axonal growth during development and for retrograde synaptic signaling at mature synapses (2-AG)^120^. The *DAGLB* variant, rs702485 (7:6449272, r^2^= 0.306 and D’=l with rs2303361) has been previously associated with high-density lipoprotein cholesterol (HDL) previously. Pathway analysis indicate a role in the triglyceride lipase activity pathway ^121^. *DAGLB* is tolerant of synonymous mutations, but intolerant of missense and loss of function mutations (ExAC Constraint scores: synonymous=−0.76, missense=1.07, LoF=0.94, **Supplementary Data 10**). Last, this variant is not predicted to be damaging by any of the algorithms tested.

***MLXIPL*** (7:73012042, rs35332062 and 7:73020337, rs3812316, known locus) The major alleles of two common non-synonymous variants (A358 V, MAF=12%; Q241H, MAF=12%) in *MLXIPL* (MLX interacting protein like) are associated with increased WHRadjBMI (β (SE)= 0.02 (0.0033), P-value=1.78×l0^−9^; β (SE)= 0.0213 (0.0034), P-value=1.98×10^−10^). These variants are in strong linkage disequilibrium (r^2^=1.00, D’=1.00, 1000 Genomes CEU). This gene encodes a basic helix-loop-helix leucine zipper transcription factor of the Myc/Max/Mad superfamily. This protein forms a heterodimeric complex and binds and activates carbohydrate response element (ChoRE) motifs in the promoters of triglyceride synthesis genes in a glucose-dependent manner^74,75^. This gene is possibly involved in the growth hormone signaling pathway and lipid metabolism. The WHRadjBMI-associated variant rs3812316 in this gene has been associated with the risk of non-alcoholic fatty liver disease and coronary artery disease^74,122,123^. Furthermore, Williams-Beuren syndrome (an autosomal dominant disorder characterized by short stature, abnormal weight gain, various cardiovascular defects, and mental retardation) is caused by a deletion of about 26 genes from the long arm of chromosome 7 including *MLXIPL*. *MLXIPL* is generally intolerant to variation, and therefore conserved (ExAC Constraint scores: synonymous = 0.48, missense=1.16, LoF=0.68, **Supplementary Data 10**). Last, both variants reported here are predicted as possible or probably damaging by one of the algorithms tested (PolyPhen).

***RAPGEF3*** (12:48143315, rsl45878042, novel locus) The major allele of a low frequency non-synonymous variant in Rap Guanine-Nucleotide-Exchange Factor (GEF) 3 (rsl45878042, L300P, MAF=1.1%) is associated with increased WHRadjBMI (β (SE)=0.085 (0.010), P-value = 7.15E^−17^). *RAPGEF3* codes for an intracellular cAMP sensor, also known as Epac (the Exchange Protein directly Activated by Cyclic AMP). Among its many known functions, RAPGEF3 regulates the ATP sensitivity of the KATP channel involved in insulin secretion^124^, may be important in regulating adipocyte differentiation^125–127^, plays an important role in regulating adiposity and energy balance^128^. *RAPGEF3* is tolerant of mutations (ExAC Constraint Scores: synonymous = −0.47, nonsynonymous = 0.32, LoF = 0, **Supplementary Data 10**). Last, our lead variant is predicted as damaging or possibly damaging for all five algorithms examined (SIFT, Polyphen2/HDIV, Polyphen2/HVAR, LRT, MutationTaster).

***TBX15*** (1:119427467, rs61730011, known locus) The major allele of a low frequency non-synonymous variant in T-box 15 (rs61730011, M460R, MAF=4.3%) is associated with increased WHRadjBMI (β(SE)=0.041(0.005)). T-box 15 (*TBX15*) is a developmental transcription factor expressed in adipose tissue, but with higher expression in visceral adipose tissue than in subcutaneous adipose tissue, and is strongly downregulated in overweight and obese individuals^129^. *TBX15* negatively controls depot-specific adipocyte differentiation and function^130^ and regulates glycolytic myofiber identity and muscle metabolism^131^. *TBX15* is moderately intolerant of mutations and therefore conserved (ExAC Constraint Scores: synonymous = 0.42, nonsynonymous = 0.65, LoF = 0.88, **Supplementary Data 10**). Last, our lead variant is predicted as damaging or possibly damaging for four of five algorithms (Polyphen2/HDIV, Polyphen2/HVAR, LRT, MutationTaster).

Overall, we observe fewer significant associations between WHRadjBMI and coding variants on the ExomeChip than Turcot *et al.*^25^ examining the association of low frequency and rare coding variants with BMI. In line with these observations, we identify fewer pathways and cross-trait associations. One reason for fewer WHRadjBMI implicated variants and pathways may be smaller sample size (N_WHRadjBMI_=476,546, N_BMI_=718,639), and thus, lower statistical power. Power, however, is likely not the only contributing factor. For example, Turcot *et al.*^25^ have comparative sample sizes between BMI and that of Marouli *et al.*^22^ studying height (N_height_=711,428). However, greater than seven times the number of coding variants are identified for height than for BMI, indicating that perhaps a number of other factors, including trait architecture, heritability (possibly overestimated in some phenotypes), and phenotype precision, likely all contribute to our study’s capacity to identify low frequency and rare variants with large effects. Further, it is possible that the comparative lack of significant findings for WHRadjBMI and BMI compared to height may be a result of higher selective pressure against genetic predisposition to cardiometabolic phenotypes, such as BMI and WHR. As evolutionary theory predicts that harmful alleles will be low frequency^61^, we may need larger sample sizes to detect rare variants that have so far escaped selective pressures. Lastly, the ExomeChip is limited by the variants that are present on the chip, which was largely dictated by sequencing studies in European-ancestry populations and a MAF detection criteria of ~0.012%. It is likely that through an increased sample size, use of chips designed to detect variation across a range of continental ancestries, high quality, deep imputation with large reference samples (e.g. HRC), and/or alternative study designs, future studies will detect additional variation from the entire allele frequency spectrum that contributes to fat distribution phenotypes.

The collected genetic and epidemiologic evidence has now demonstrated that fat distribution (as measured by increased WHRadjBMI) is correlated with increased risk of T2D and CVD, and that this association is likely causal with potential mediation through blood pressure, triglyceride-rich lipoproteins, glucose, and insulin^9^. This observation yields an immediate follow-up question: Which mechanisms regulate depot-specific fat accumulation and are risks for disease, driven by increased visceral or decreased subcutaneous adipose tissue mass (or both)? Pathway analysis identified several novel pathways and gene sets related to metabolism and adipose regulation, bone growth and development we also observed a possible role for adiponectin, a hormone which has been linked to “healthy” expansion of adipose tissue and insulin sensitivity^62^. Similarly, expression/eQTL results support the function and relevance of adipogenesis, adipocyte biology, and insulin signaling, supporting our previous findings for WHRadjBMI^10^. We also provide evidence suggesting known biological functions and pathways contributing to body fat distribution (e.g., diet-induced obesity, angiogenesis, bone growth and morphology, and enhanced lipolysis).

The ultimate aim of genetic investigations of obesity-related traits, like those presented here, is to identify genomic pathways that are dysregulated leading to obesity pathogenesis, and may result in a myriad of downstream illnesses. Thus, our findings may enhance the understanding of central obesity and identify new molecular targets to avert its negative health consequences. Significant cross-trait associations and additional associations observed in the GWAS Catalog are consistent with expected direction of effect for several traits, i.e. the WHR-increasing allele is associated with higher values of TG, DBP, fasting insulin, TC, LDL and T2D across many significant variants. However, it is worth noting that there are some exceptions. For example, rs9469913-A in *UHRF1BP1* is associated with both increased WHRadjBMI and increased HDL. Also, we identified two variants in *MLXIPL* (rs3812316 and rs35332062), a well-known lipids-associated locus, in which the WHRadjBMI-increasing allele also increases all lipid levels, risk for hypertriglyceridemia, SBP and DBP. However, our findings show a significant and negative association with HbA1C, and nominally significant and negative associations with two-hour glucose, fasting glucose, and Type 2 diabetes, and potential negative associations with biomarkers for liver disease (e.g. gamma glutamyl transpeptidase). Other notable exceptions include *ITIH3* (negatively associated with BMI, HbA1C, LDL and SBP), *DAGLB* (positively associated with HDL), and *STAB1* (negatively associated with TC, LDL, and SBP in cross-trait associations). Therefore, caution in selecting pathways for therapeutic targets is warranted; one must look beyond the effects on central adiposity, but also at the potential cascading effects of related diseases.

A seminal finding from this study is the importance of lipid metabolism for body fat distribution. In fact, pathway analyses that highlight enhanced lipolysis, cross-trait associations with circulating lipid levels, existing biological evidence from the literature, and knockdown experiments in *Drosophila* examining triglyceride storage point to novel candidate genes (*ANGPTL4, ACVR1C, DAGLB, MGA, RASIP1*, and *IZUMO1*) and new candidates in known regions (*DNAH10*^10^ and *MLXIPL*^14^) related to lipid biology and its role in fat storage. Newly implicated genes of interest include *ACVR1C, MLXIPL*, and *ANGPTL4*, all of which are involved in lipid homeostasis; all are excellent candidate genes for central adiposity. Carriers of inactivating mutations in *ANGPTL4 {Angiopoietin Like 4*), for example, display low triglyceride levels and low risk of coronary artery disease^63^. *ACVR1C* encodes the activin receptor-like kinase 7 protein (ALK7), a receptor for the transcription factor TGFB-1, well known for its central role in growth and development in general^64–68^, and adipocyte development in particular^68^. *ACVR1C* exhibits the highest expression in adipose tissue, but is also highly expressed in the brain^69–71^. In mice, decreased activity of *ACVR1C* upregulates PPARγ and C/EBPα pathways and increases lipolysis in adipocytes, thus decreasing weight and diabetes in mice^69,72,73^. Such activity is suggestive of a role for ALK7 in adipose tissue signaling and therefore for therapeutic targets for human obesity. *MLXIPL*, also important for lipid metabolism and postnatal cellular growth, is a transcription factor which activates triglyceride synthesis genes in a glucose-dependent manner^74,75^. The lead exome variant in this gene is highly conserved, most likely damaging, and is associated with reduced *MLXIPL* expression in adipose tissue. Furthermore, in a recent longitudinal, *in vitro* transcriptome analysis of adipogenesis in human adipose-derived stromal cells, gene expression of *MLXIPL* was up-regulated during the maturation of adipocytes, suggesting a critical role in the regulation of adipocyte size and accumulation^76^. However, given our observations on cross-trait associations with variants in *MLXIPL* and diabetes-related traits, development of therapeutic targets must be approached cautiously.

Taken together, our 24 novel variants for WHRadjBMI offer new biology, highlighting the importance of lipid metabolism in the genetic underpinnings of body fat distribution. We continue to demonstrate the critical role of adipocyte biology and insulin resistance for central obesity and offer support for potentially causal genes underlying previously identified fat distribution GWAS loci. Notably, our findings offer potential new therapeutic targets for intervention in the risks associated with abdominal fat accumulation, and represents a major advance in our understanding of the underlying biology and genetic architecture of central adiposity.

## ACKNOWLEDGEMENTS

A full list of acknowledgements is provided in the **Supplementary Table 17**. This study was completed as part of the Genetic Investigation of ANtropometric Traits (GIANT) Consortium. This research has been conducted using the UK Biobank resource. Funding for this project was provided by Aase and Ejner Danielsens Foundation, Academy of Finland (102318; 123885; 117844; 40758; 211497; 118590; 139635; 129293; 286284; 134309; 126925; 121584; 124282; 129378; 117787; 41071; 137544; 272741), Action on Hearing Loss (G51), ALK-Abelló A/S (Hørsholm-Denmark), American Heart Association (13EIA14220013; 13GRNT16490017; 13POST16500011), American Recovery and Reinvestment Act of 2009 (ARRA) Supplement (EY014684-03S1; −04S1), Amgen, André and France Desmarais Montreal Heart Institute (MHI) Foundation, AstraZeneca, Augustinus Foundation, Australian Government and Government of Western Australia, Australian Research Council Future Fellowship, Becket Foundation, Benzon Foundation, Bernard Wolfe Health Neuroscience Endowment, British Heart Foundation (CH/03/001; RG/14/5/30893; RG/200004; SP/04/002; SP/09/002), BiomarCaRE (278913), Bundesministerium für Bildung und Forschung (Federal Ministry of Education and Research-Germany; German Center for Diabetes Research (DZD); 01ER1206; 01ER1507; 01ER1206; 01ER1507; FKZ: 01E01501 (AD2-060E); 01ZZ9603; 01ZZ0103; 01ZZ0403; 03IS2061A; 03Z1CN22; FKZ 01GI1128), Boehringer Ingelheim Foundation, Boston University School of Medicine, Canada Research Chair program, Canadian Cancer Society Research Institute, Canadian Institutes of Health Research (MOP-82893), Cancer Research UK (C864/A14136; A490/A10124; C8197/A16565), Cebu Longitudinal Health and Nutrition Survey (CLHNS) pilot funds (RR020649; ES010126; DK056350), Center for Non-Communicable Diseases (Pakistan), Central Society for Clinical Research, Centre National de Génotypage (Paris-France), CHDI Foundation (Princeton-USA), Chief Scientist Office of the Scottish Government Health Directorate (CZD/16/6), City of Kuopio and Social Insurance Institution of Finland (4/26/2010), Clarendon Scholarship, Commission of the European Communities; Directorate C-Public Health (2004310), Copenhagen County, County Council of Dalarna, Curtin University of Technology, Dalarna University, Danish Centre for Evaluation and Health Technology Assessment, Danish Council for Independent Research, Danish Diabetes Academy, Danish Heart Foundation, Danish Medical Research Council-Danish Agency for Science Technology and Innovation, Danish Medical Research Council, Danish Pharmaceutical Association, Danish Research Council for Independent Research, Dekker scholarship (2014T001), Dentistry and Health Sciences, Department of Internal Medicine at the University of Michigan, Diabetes Care System West-Friesland, Diabetes Heart Study (ROI HL6734; ROI HL092301; R01 NS058700), Doris Duke Charitable Foundation Clinical Scientist Development Award (2014105), Doris Duke Medical Foundation, Dr. Robert Pfleger Stiftung, Dutch Cancer Society (NKI2009-4363), Dutch Government (NWO 184.021.00; NWO/MaGW VIDI-016-065-318; NWO VICI 453-14-0057; NWO 184.021.007), Dutch Science Organization (ZonMW-VENI Grant 916.14.023), Edith Cowan University, Education and Sports Research Grant (216-1080315-0302); Croatian Science Foundation (grant 8875), Else Kröner-Frsenius-Stiftung (2012_A147), Emil Aaltonen Foundation, Erasmus Medical Center, Erasmus University (Rotterdam), European Research Council Advanced Principal Investigator Award, European Research Council (310644; 268834; 323195; SZ-245 50371-GLUCOSEGENES-FP7-IDEAS-ERC; 293574), Estonian Research Council (IUT20-60), European Union Framework Programme 6 (LSHM_CT_2006_037197; Bloodomics Integrated Project; LSHM-CT-2004-005272; LSHG-CT-2006-018947), European Union Framework Programme 7 (HEALTH-F2-2013-601456; HEALTH-F2-2012-279233; 279153; HEALTH-F3-2010-242244; EpiMigrant; 279143; 313010; 305280; HZ2020 633589; 313010; HEALTH-F2-2011-278913; HEALTH-F4-2007-201413), European Commission (DG XII), European Community (SOC 98200769 05 F02), European Regional Development Fund to the Centre of Excellence in Genomics and Translational Medicine (GenTransMed), European Union (QLG1-CT-2001-01252; SOC 95201408 05 F02), EVO funding of the Kuopio University Hospital from Ministry of Health and Social Affairs (5254), Eye Birth Defects Foundation Inc., Federal Ministry of Science-Germany (01 EA 9401), Finland’s Slottery Machine Association, Finnish Academy (255935; 269517), Finnish Cardiovascular Research Foundation, Finnish Cultural Foundation, Finnish Diabetes Association, Finnish Diabetes Research Foundation, Finnish Foundation for Cardiovascular Research, Finnish Funding Agency for Technology and Innovation (40058/07), Finnish Heart Association, Finnish National Public Health Institute, Fondation Leducq (14CVD01), Food Standards Agency (UK), Framingham Heart Study of the National Heart Lung and Blood Institute of the National Institutes of Health (HHSN268201500001; N02-HL-6-4278), FUSION Study (DK093757; DK072193; DK062370; ZIA-HG000024), General Clinical Research Centre of the Wake Forest School of Medicine (MOI RR07122; F32 HL085989), Genetic Laboratory of the Department of Internal Medicine-Erasmus MC (the Netherlands Genomics Initiative), Genetics and Epidemiology of Colorectal Cancer Consortium (NCI CA137088), German Cancer Aid (70-2488-Ha I), German Diabetes Association, German Research Foundation (CRC 1052 C01; B01; B03), Health and Retirement Study (R03 AG046398), Health Insurance Foundation (2010 Β 131), Health Ministry of Lombardia Region (Italy), Helmholtz Zentrum München - German Research Center for Environmental Health, Helse Vest, Home Office (780-TETRA), Hospital Districts of Pirkanmaa; Southern Ostrobothnia; North Ostrobothnia; Central Finland and Northern Savo, Ib Henriksen Foundation, Imperial College Biomedical Research Centre, Imperial College Healthcare NHS Trust, Institute of Cancer Research and The Everyman Campaign, Interuniversity Cardiology Institute of the Netherlands (09.001), Intramural Research Program of the National Institute on Aging, Italian Ministry of Health (GR-2011-02349604), Johns Hopkins University School of Medicine (HHSN268200900041C), Juho Vainio Foundation, Kaiser Foundation Research Institute (HHSN268201300029C), KfH Stiftung Präventivmedizin e.V., KG Jebsen Foundation, Knut and Alice Wallenberg Foundation (Wallenberg Academy Fellow), Knut och Alice Wallenberg Foundation (2013.0126), Kuopio Tampere and Turku University Hospital Medical Funds (X51001), Kuopio University Hospital, Leenaards Foundation, Leiden University Medical Center, Li Ka Shing Foundation (CML), Ludwig-Maximilians-Universität, Lund University, Lundbeck Foundation, Lundbeckfonden, Marianne and Marcus Wallenberg Foundation, Max Planck Society, Medical Research Council-UK (G0601966; G0700931; G0000934; MR/L01632X/1; MC_UU_12015/1; MC_PC_13048; G9521010D; G1000143; MC_UU_12013/l-9; MC_UU_12015/1; MC_PC_13046; MC_U106179471; G0800270, MR/L01341X/1), MEKOS Laboratories (Denmark), Merck & Co Inc., MESA Family (R01-HL-071205; R01-HL-071051; R01-HL-071250; R01-HL-071251; R01-HL-071252; R01-HL-071258; R01-HL-071259; UL1-RR-025005), Ministry for Health Welfare and Sports (the Netherlands), Ministry of Cultural Affairs (Germany), Ministry of Education and Culture of Finland (627;2004-2011), Ministry of Education Culture and Science (the Netherlands), Ministry of Science and Technology (Taiwan) (MOST 104-2314-B-075A-006 -MY3), Ministry of Social Affairs and Health in Finland, Montreal Heart Institute Foundation, MRC-PHE Centre for Environment and Health, Multi-Ethnic Study of Atherosclerosis (MESA) (N01-HC-95159; N01-HC-95160; N01-HC-95161; N01-HC-95162; N01-HC-95163; N01-HC-95164; N01-HC-95165; N01-HC-95166; N01-HC-95167; N01-HC-95168; N01-HC-95169), Munich Center of Health Sciences (MC-Health), Municipality of Rotterdam (the Netherlands) Murdoch University, National Basic Research Program of China (973 Program 2012CB524900), National Cancer Institute (CA047988; UM1CA182913), National Cancer Research Institute UK, National Cancer Research Network UK, National Center for Advancing Translational Sciences (UL1TR001881), National Center for Research Resources (UL1-TR-000040 and UL1-RR-025005), National Eye Institute of the National Institutes of Health (EY014684, EY-017337), National Health and Medical Research Council of Australia (403981; 1021105; 572613), National Heart Lung and Blood Institute (HHSN268201100010C; HHSN268201100011C; and HHSN268201100012C; HHSN268201100005; HHSN268201100006C; HHSN268201100007C; HHSN268201100008C; HHSN268201100009; HHSN268201100037C; HHSN268201300046C; HHSN268201300047C; HHSN268201300048C; HHSN268201300049C; HHSN268201300050C; HL043851; HL080467; HL094535; HL109946; HHSN268201300025C; HHSN268201300026C; HL119443; HL054464; HL054457; HL054481; HL087660; HL086694; HL060944; HL061019; HL060919; HL060944; HL061019; N02-HL-6-4278; R21 HL121422-02; R21 HL121422-02; ROI DK089256-05), National Human Genome Research Institute (HG007112), National Institute for Health Research BioResource Clinical Research Facility and Biomedical Research Centre based at Guy’s and St Thomas’ NHS Foundation Trust and King’s College London, National Institute for Health Research Comprehensive Biomedical Research Centre Imperial College Healthcare NHS Trust, National Institute for Health Research (NIHR) (RP-PG-0407-10371), National Institute of Diabetes and Digestive and Kidney Disease (DK063491; DK097524; DK085175; DK087914; 1R01DK8925601; 1R01DK106236-01A1), National Institute of Health Research Senior Investigator, National Institute on Aging (NIA U01AG009740; RC2 AG036495; RC4 AG039029), National Institute on Minority Health and Health Disparities, National Institutes of Health (NIH) (1R01HG008983-01; 1R21DA040177-01; 1R01HL092577; R01HL128914; K24HL105780; K01HL116770; U01 HL072515-06; U01 HL84756; U01HL105198; U01 GM074518; ROI DK089256-05; R01DK075787; R25 CA94880; P30 CA008748; DK078150; TW005596; HL085144; TW008288; R01-HL093029; U01-HG004729; R01-DK089256; 1R01DK101855-01; 1K99HL130580; T32-GM067553; U01-DK105561; R01-HL-117078; R01-DK-089256; U01HG008657; U01HG06375; U01AG006781; DK064265; R01DK106621-01; K23HL114724; NS33335; HL57818; R01-DK089256; 2R01HD057194; U01HG007416; R01DK101855, R01DK075787, T32 GM096911-05; KOI DK107836; R01DK075787; UOl AG 06781; U01-HG005152, 1F31HG009850-01), National Key R&D Plan of China (2016YFC1304903), Key Project of the Chinese Academy of Sciences (ZDBS-SSW-DQC-02, ZDRW-ZS-2016-8-1, KJZD-EW-L14-2-2), National Natural Science Foundation of China (81471013; 30930081; 81170734; 81321062; 81471013), National NIHR Bioresource, National Science Council (Taiwan) (NSC 102-2314-B-075A-002), Netherlands Cardiovascular Research Initiative (CVON2011-19), Netherlands Heart Foundation, Netherlands Organisation for Health Research and Development (ZonMW) (113102006), Netherlands Organisation for Scientific Research (NWO)-sponsored Netherlands Consortium for Healthy Aging (050-060-810), Netherlands Organization for Scientific Research (184021007), NHMRC Practitioner Fellowship (APP1103329), NIH through the American Recovery and Reinvestment Act of 2009 (ARRA) (5RC2HL102419), NIHR Biomedical Research Centre at The Institute of Cancer Research and The Royal Marsden NHS Foundation Trust, NIHR Cambridge Biomedical Research Centre, NIHR Cambridge Biomedical Research Centre, NIHR Health Protection Research Unit on Health Impact of Environmental Hazards (HPRU-2012-10141), NIHR Leicester Cardiovascular Biomedical Research Unit, NIHR Official Development Assistance (ODA, award 16/136/68), NIHR Oxford Biomedical Research Centre, the European Union FP7 (EpiMigrant, 279143) and H2020 programs (iHealth-T2D; 643774), NIHR Senior Investigator, Nordic Centre of Excellence on Systems Biology in Controlled Dietary Interventions and Cohort Studies (SYSDIET) (070014), Northwestern University (HHSN268201300027C), Norwegian Diabetes Association, Novartis, Novo Nordisk Foundation, Nuffield Department of Clinical Medicine Award, Orchid Cancer Appeal, Oxford Biomedical Research Centre, Paavo Nurmi Foundation, Päivikki and Sakari Sohlberg Foundation, Pawsey Supercomputing Centre (funded by Australian Government and Government of Western Australia), Peninsula Research Bank-NIHR Exeter Clinical Research Facility, Pfizer, Prostate Cancer Research Foundation, Prostate Research Campaign UK (now Prostate Action), Public Health England, QIMR Berghofer, Raine Medical Research Foundation, Regione FVG (L.26.2008), Republic of Croatia Ministry of Science, Research Centre for Prevention and Health-the Capital Region of Denmark, Research Council of Norway, Research Institute for Diseases in the Elderly (RIDE), Research into Ageing, Robert Dawson Evans Endowment of the Department of Medicine at Boston University School of Medicine and Boston Medical Center, Science Live/Science Center NEMO, Scottish Funding Council (HR03006), Sigrid Juselius Foundation, Social Insurance Institution of Finland, Singapore Ministry of Health’s National Medical Research Council (NMRC/STaR/0028/2017), Social Ministry of the Federal State of Mecklenburg-West Pomerania, State of Bavaria-Germany, State of Washington Life Sciences Discovery Award (265508) to the Northwest Institute of Genetic Medicine, Stroke Association, Swedish Diabetes Foundation (2013­024), Swedish Heart-Lung Foundation (20120197; 20120197; 20140422), Swedish Research Council (2012-1397), Swedish Research Council Strategic Research Network Epidemiology for Health, Swiss National Science Foundation (31003A-143914), SystemsX.ch (51RTP0_151019), Taichung Veterans General Hospital (Taiwan) (TCVGH-1047319D; TCVGH-1047311C), Tampere Tuberculosis Foundation, TEKES Grants (70103/06; 40058/07), The Telethon Kids Institute, Timber Merchant Vilhelm Bangs Foundation, UCL Hospitals NIHR Biomedical Research Centre, UK Department of Health, Université de Montréal Beaulieu-Saucier Chair in Pharmacogenomics, University Hospital Regensburg, University of Bergen, University of Cambridge, University of Michigan Biological Sciences Scholars Program, University of Michigan Internal Medicine Department Division of Gastroenterology, University of Minnesota (HHSN268201300028C), University of Notre Dame (Australia), University of Queensland, University of Western Australia (UWA), Uppsala Multidisciplinary Center for Advanced Computational Science (b2011036), Uppsala University, US Department of Health and Human Services (HHSN268201100046C; HHSN268201100001C; HHSN268201100002C; HHSN268201100003C; HHSN268201100004C; HHSN271201100004C), UWA Faculty of Medicine, Velux Foundation, Wellcome Trust (083948/B/07/Z; 084723/Z/08/Z; 090532; 098381; 098497/Z/12/Z; WT098051; 068545/Z/02; WT064890; WT086596; WT098017; WT090532; WT098051; WT098017; WT098381; WT098395; 083948; 085475), Western Australian DNA Bank (National Health and Medical Research Council of Australia National Enabling Facility), Women and Infant’s Research Foundation, Yrjš Jahnsson Foundation (56358)

## AUTHORSHIP CONTRIBUTIONS

Writing Group: LAC, RSF, TMF, MG, HMH, JNH, AEJ, TK, ZK, CML, RJFL, YL, KEN, VT, KLY; Data preparation group: TA, IBB, TE, SF, MG, HMH, AEJ, TK, DJL, KSL, AEL, RJFL, YL, EM, NGDM, MCMG, PM, MCYN, MAR, SS, CS, KS, VT, SV, SMW, TWW, KLY, XZ; WHR meta-analyses: PLA, HMH, AEJ, TK, MG, CML, RJFL, KEN, VT, KLY; Pleiotropy working group: GA, MB, JPC, PD, FD, JCF, HMH, SK, HK, HMH, AEJ, CML, DJL, RJFL, AM, EM, GM, MIM, PBM, GMP, JRBP, KSR, XS, SW, JW, CJW; Phenome-wide association studies: LB, JCD, TLE, AG, AM, MIM; Gene-set enrichment analyses: SB, RSF, JNH, ZK, DL, THP; eQTL analyses: CKR, YL, KLM; Monogenic and syndromic gene enrichment analyses: HMH, AKM; Fly Obesity Screen: AL, JAP; Overseeing of contributing studies: (1958 Birth Cohort) PD; (Airwave) PE; (AMC PAS) GKH; (Amish) JRO’C; (ARIC) EB; (ARIC, Add Health) KEN; (BRAVE) EDA, RC; (BRIGHT) PBM; (CARDIA) MF, PJS; (Cebu Longitudinal Health and Nutrition Survey) KLM; (CHD Exome + Consortium) ASB, JMMH, DFR, JD; (CHES) RV; (Clear/eMERGE (Seattle)) GPJ; (CROATIA_Korcula) VV, OP, IR; (deCODE) KS, UT; (DHS) DWB; (DIACORE) CAB; (DPS) JT, JL, MU; (DRSEXTRA) TAL, RR; (EFSOCH) ATH, TMF; (EGCUT) TE; (eMERGE (Seattle)) EBL; (EPIC-Potsdam) MBS, HB; (EpiHealth) EI, PWF; (EXTEND) ATH, TMF; (Family Heart Study) IBB; (Fenland, EPIC) RAS; (Fenland, EPIC, InterAct) NJW, CL; (FINRISK) SM; (FINRISK 2007 (T2D)) PJ, VS; (Framingham Heart Study) LAC; (FUSION) MB, FSC; (FVG) PG; (Generation Scotland) CH, BHS; (Genetic Epidemiology Network of Arteriopathy (GENOA)) SLRK; (GRAPHIC) NJS; (GSK-STABILITY) DMW, LW, HDW; (Health) AL; (HELIC MANOLIS) EZ, GD; (HELIC Pomak) EZ, GD; (HUNT-MI) KH, CJW; (Inter99) TH, TJ; (IRASFS) LEW, EKS; (Jackson Heart Study (JHS)) JGW; (KORA S4) KS, IMH; (Leipzig-Adults) MB, PK; (LOLIPOP-Exome) JCC, JSK; (LOLIPOP-OmniEE) JCC, JSK; (MESA) JIR, XG; (METSIM) JK, ML; (MONICA-Brianza) GC; (Montreal Heart Institute Biobank (MHIBB)) MPD, GL, SdD, JCT; (MORGAM Central Laboratory) MP; (MORGAM Data Centre) KK; (OBB) FK; (PCOS) APM, CML; (PIVUS) CML, LL; (PRIME - Belfast) FK; (PRIME - Lille) PA; (PRIME - Strasbourg) MM; (PRIME - Toulouse) JF; (PROMIS) DS; (QC) MAR; (RISC) BB, EF, MW; (Rotterdam Study I) AGU, MAI; (SEARCH) AMD; (SHIP/SHIP-Trend) MD; (SIBS) DFE; (SOLID TIMI-52) DMW; (SORBS) APM, MS, AT; (The Mount Sinai BioMe Biobank) EPB, RJFL; (The NEO Study) DOMK; (The NHAPC study, The GBTDS study) XL; (The Western Australian Pregnancy Cohort (Raine) Study) CEP, SM; (TwinsUK) TDS; (ULSAM) APM; (Vejle Biobank) IB, CC, OP; (WGHS) DIC, PMR; (Women’s Health Initiative) PLA; (WTCCC-UKT2D) MIM, KRO; (YFS) TL, OTRa; Genotyping of contributing studies: (1958 Birth Cohort) KES; (Airwave) EE, MPSL; (AMC PAS) SS; (Amish) LMYA, JAP; (ARIC) EWD, MG; (BBMRI-NL) SHV, LB, CMvD, PIWdB; (BRAVE) EDA; (Cambridge Cancer Studies) JGD; (CARDIA) MF; (CHD Exome + Consortium) ASB, JMMH, DFR, JD, RY(Clear/eMERGE (Seattle)) GPJ; (CROATIA_Korcula) VV; (DIACORE) CAB, MG; (DPS) AUJ, JL; (DRSEXTRA) PK; (EGCUT) TE; (EPIC-Potsdam) MBS, KM; (EpiHealth) EI, PWF; (Family Heart Study) KDT; (Fenland, EPIC) RAS; (Fenland, EPIC, InterAct) NJW, CL; (FUSION) NN; (FVG) IG, AM; (Generation Scotland) CH; (Genetic Epidemiology Network of Arteriopathy (GENOA)) SLRK, JAS; (GRAPHIC) NJS; (GSK-STABILITY) DMW; (Health) JBJ; (HELIC MANOLIS) LS; (HELIC Pomak) LS; (Inter99) TH, NG; (KORA) MMN; (KORA S4) KS, HG; (Leipzig-Adults) AM; (LOLIPOP-Exome) JCC, JSK; (LOLIPOP-OmniEE) JCC, JSK; (MESA) JIR, YDIC, KDT; (METSIM) JK, ML; (Montreal Heart Institute Biobank (MHIBB)) MPD; (OBB) FK; (PCOS) APM; (PIVUS) CML; (Rotterdam Study I) AGU, CMG, FR; (SDC) JMJ, HV; (SEARCH) AIMD; (SOLID TIMI-52) DMW; (SORBS) APM; (The Mount Sinai BioMe Biobank) EPB, RJFL, YL, CS; (The NEO Study) RLG; (The NHAPC study, The GBTDS study) XL, HL, YH; (The Western Australian Pregnancy Cohort (Raine) Study) CEP, SM; (TUDR) ZA; (TwinsUK) APM; (ULSAM) APM; (WGHS) DIC, AYC; (Women’s Health Initiative) APR; (WTCCC-UKT2D) MIM; (YFS) TL, LPL; Phenotyping of contributing studies: (Airwave) EE; (AMC PAS) SS; (Amish) LM YA; (ARIC) EWD; (ARIC, Add Health) KEN; (BBMRI-NL) SHV; (BRAVE) EDA; (BRIGHT) MJC; (CARL) AR, GG; (Cebu Longitudinal Health and Nutrition Survey) NRL; (CHES) RV, MT; (Clear/eMERGE (Seattle)) GPJ, AAB; (CROATIA_Korcula) OP, IR; (DIACORE) CAB, BKK; (DPS) AUJ, JL; (EFSOCH) ATH; (EGCUT) EM; (EPIC-Potsdam) HB; (EpiHealth) EI; (EXTEND) ATH; (Family Heart Study) MFF; (Fenland, EPIC, InterAct) NJW; (FIN-D2D 2007) LM, MV; (FINRISK) SM; (FINRISK 2007 (T2D)) PJ, HS; (Framingham Heart Study) CSF; (Generation Scotland) CH, BHS; (Genetic Epidemiology Network of Arteriopathy (GENOA)) SLRK, JAS; (GRAPHIC) NJS; (GSK-STABILITY) LW, HDW; (Health) AL, BHT; (HELIC MANOLIS) LS, AEF, ET; (HELIC Pomak) LS, AEF, MK; (HUNT-MI) KH, OH; (Inter99) TJ, NG; (IRASFS) LEW, BK; (KORA) MMN; (LASA (BBMRI-NL)) KM AS; (Leipzig-Adults) MB, PK; (LOLIPOP-Exome) JCC, JSK; (LOLIPOP-OmniEE) JCC, JSK; (MESA) MA; (Montreal Heart Institute Biobank (MHIBB)) GL, KSL, VT; (MORGAM Data Centre) KK; (OBB) FK, MN; (PCOS) CML; (PIVUS) LL; (PRIME - Belfast) FK; (PRIME - Lille) PA; (PRIME - Strasbourg) MM; (PRIME - Toulouse) JF; (RISC) BB, EF; (Rotterdam Study I) MAI, CMGFR, MCZ; (SHIP/SHIP-Trend) NF; (SORBS) MS, AT; (The Mount Sinai BioMe Biobank) EPB, YL, CS; (The NEO Study) RdM; (The NHAPC study, The GBTDS study) XL, HL, LS, FW; (The Western Australian Pregnancy Cohort (Raine) Study) CEP; (TUDR) YJH, WJL; (TwinsUK) TDS, KSS; (ULSAM) VG; (WGHS) DIC, PMR; (Women’s Health Initiative) APR; (WTCCC-UKT2D) MIM, KRO; (YFS) TL, OTR; Data analysis of contributing studies: (1958 Birth Cohort) KES, IN; (Airwave) EE, MPSL; (AMC PAS) SS; (Amish) JRO’C, LMYA, JAP; (ARIC, Add Health) KEN, KLY, MG; (BBMRI-NL) LB; (BRAVE) RC, DSA; (BRIGHT) HRW; (Cambridge Cancer Studies) JGD, AP, DJT; (CARDIA) MF, LAL; (CARL) AR, DV; (Cebu Longitudinal Health and Nutrition Survey) YW; (CHD Exome + Consortium) ASB, JMMH, DFR, RY, PS; (CHES) YJ; (CROATIA_Korcula) VV; (deCODE) VS, GT; (DHS) AJC, PM, MCYN; (DIACORE) CAB, MG; (EFSOCH) HY; (EGCUT) TE, RM; (eMERGE (Seattle)) DSC; (ENDO) TK; (EPIC) JHZ; (EPIC-Potsdam) KM; (EpiHealth) SG; (EXTEND) HY; (Family Heart Study) MFF; (Fenland) JaL; (Fenland, EPIC) RAS; (Fenland, InterAct) SMW; (Finrisk Extremes and QC) SV; (Framingham Heart Study) CTL, NLHC; (FVG) IG; (Generation Scotland) CH, JM; (Genetic Epidemiology Network of Arteriopathy (GENOA)) LFB; (GIANT-Analyst) AEJ; (GRAPHIC) NJS, NGDM, CPN; (GSK-STABILITY) DMW, AS; (Health) JBJ; (HELIC MANOLIS) LS; (HELIC Pomak) LS; (HUNT-MI) WZ; (Inter99) NG; (IRASFS) BK; (Jackson Heart Study (JHS)) LAL, JL; (KORA S4) TWW; (LASA (BBMRI-NL)) KMAS; (Leipzig-Adults) AM; (LOLIPOP-Exome) JCC, JSK, WZ; (LOLIPOP-OmniEE) JCC, JSK, WZ; (MESA) JIR, XG, JY; (METSIM) XS; (Montreal Heart Institute Biobank (MHIBB)) JCT, GL, KSL, VT; (OBB) AM; (PCOS) APM, TK; (PIVUS) NR; (PROMIS) AR, WZ; (QC G0T2D/T2 D-GENES (FUSION, METSIM, etc)) AEL; (RISC) HY; (Rotterdam Study I) CMG, FR; (SHIP/SHIP-Trend) AT; (SOLID TIMI-52) DMW, AS; (SORBS) APM; (The Mount Sinai BioMe Biobank) YL, CS; (The NEO Study) RLG; (The NHAPC study, The GBTDS study) XL, HL, YH; (The Western Australian Pregnancy Cohort (Raine) Study) CAW; (UK Biobank) ARW; (ULSAM) APM, AM; (WGHS) DIC, AYC; (Women’s Health Initiative) PLA, JH; (WTCCC-UKT2D) WG; (YFS) LPL.

## METHODS

### Studies

Stage 1 consisted of 74 studies (12 case/control studies, 59 population-based studies, and five family studies) comprising 344,369 adult individuals of the following ancestries: 1) European descent (N=288,492), 2) African (N=15,687), 3) South Asian (N=29,315), 4) East Asian (N=6,800), and 5) Hispanic (N=4,075). Stage 1 meta-analyses were carried out in each ancestry separately and in the all ancestries group, for both sex-combined and sex-specific analyses. Follow-up analyses were undertaken in 132,177 individuals of European ancestry from the deCODE anthropometric study and UK Biobank **(Supplementary Tables 1-3)**. Conditional analyses were performed in the all ancestries and European descent groups. Informed consent was obtained for participants by the parent study and protocols approved by each study’s institutional review boards.

### Phenotypes

For each study, WHR (waist circumference divided by hip circumference) was corrected for age, BMI, and the genomic principal components (derived from GWAS data, the variants with MAF >1% on the ExomeChip, and ancestry informative markers available on the ExomeChip), as well as any additional study-specific covariates (e.g. recruiting center), in a linear regression model. For studies with non-related individuals, residuals were calculated separately by sex, whereas for family-based studies sex was included as a covariate in models with both men and women. Additionally, residuals for case/control studies were calculated separately. Finally, residuals were inverse normal transformed and used as the outcome in association analyses. Phenotype descriptives by study are shown in **Supplementary Table 3**.

### Genotypes and QC

The majority of studies followed a standardized protocol and performed genotype calling using the algorithms indicated in **Supplementary Table 2**, which typically included zCall^3^. For 10 studies participating in the Cohorts for Heart and Aging Research in Genomic Epidemiology (CHARGE) Consortium, the raw intensity data for the samples from seven genotyping centers were assembled into a single project for joint calling^4^. Study-specific quality control (QC) measures of the genotyped variants were implemented before association analysis **(Supplementary Tables 1-2)**. Furthermore, to assess the possibility that any significant associations with rare and low-frequency variants could be due to allele calling in the smaller studies, we performed a sensitivity meta-analysis including all large studies (>5,000 participants) and compared to all studies. We found very high concordance for effect sizes, suggesting that smaller studies do not bias our results **(Supplementary Fig. 24)**.

### Study-level statistical analyses

Individual cohorts were analyzed for each ancestry separately, in sex-combined and sex-specific groups, with either RAREMETALWORKER (http://genome.sph.umich.edu/wiki/RAREMETALWORKER) or RVTESTs (http://zhanxw.github.io/rvtests/), to associate inverse normal transformed WHRadjBMI with genotype accounting for cryptic relatedness (kinship matrix) in a linear mixed model. These software programs are designed to perform score-statistic based rare-variant association analysis, can accommodate both unrelated and related individuals, and provide single-variant results and variance-covariance matrices. The covariance matrix captures linkage disequilibrium (LD) relationships between markers within 1 Mb, which is used for gene-level meta-analyses and conditional analyses^77,78^. Single-variant analyses were performed for both additive and recessive models.

### Centralized quality-control

Individual cohorts identified ancestry population outliers based on 1000 Genome Project phase 1 ancestry reference populations. A centralized quality-control procedure implemented in EasyQC^79^ was applied to individual cohort association summary statistics to identify cohort-specific problems: (1) assessment of possible errors in phenotype residual transformation; (2) comparison of allele frequency alignment against 1000 Genomes Project phase 1 reference data to pinpoint any potential strand issues, and (3) examination of quantile-quantile (QQ) plots per study to identify any inflation arising from population stratification, cryptic relatedness and genotype biases.

### Meta-analyses

Meta-analyses were carried out in parallel by two different analysts at two sites using RAREMETAL^77^. During the meta-analyses, we excluded variants if they had call rate <95%, Hardy-Weinberg equilibrium P-value <1×10^−7^, or large allele frequency deviations from reference populations (>0.6 for all ancestries analyses and >0.3 for ancestry-specific population analyses). We also excluded from downstream analyses markers not present on the Illumina ExomeChip array 1.0, variants on the Y-chromosome or the mitochondrial genome, indels, multiallelic variants, and problematic variants based on the Blat-based sequence alignment analyses. Significance for single-variant analyses was defined at an array-wide level (P<2×10^−7^). For all suggestive significant variants from Stage 1, we tested for significant sex differences. We calculated Psexhet for each SNP, testing for difference between women-specific and men-specific beta estimates and standard errors using EasyStrata^11,80^. Each SNP that reached P_sexhet_<0.05/# of variants tested (70 variants brought forward from Stage 1, P_sexhet_<7.14×1O^−4^) was considered significant. Additionally, while each individual study was asked to perform association analyses stratified by race/ethnicity, and adjust for population stratification, all study-specific summary statistics were meta-analyzed together for our all ancestry meta-analyses. To investigate potential heterogeneity across ancestries, we did examine ancestry-specific meta-analysis results for our top 70 variants from stage 1, and found no evidence of significant across-ancestry heterogeneity observed for any of our top variants (l^2^ values noted in **Supplementary Data 1-3)**.

For the gene-based analyses, we applied two sets of criteria to select variants with a MAF<5% within each ancestry based on coding variant annotation from five prediction algorithms (PolyPhen2, HumDiv and HumVar, LRT, MutationTaster, and SIFT)^80,81^. Our broad gene-based tests included nonsense, stop-loss, splice site, and missense variants annotated as damaging by at least one algorithm mentioned above. Our strict gene-based tests included only nonsense, stop-loss, splice site, and missense variants annotated as damaging by all five algorithms. These analyses were performed using the sequence kernel association test (SKAT) and variable threshold (VT) methods. Statistical significance for gene-based tests was set at a Bonferroni-corrected threshold of P<2.5×10^−6^ (~0.05/>20,000 genes). All gene-based tests were performed in RAREMETAL^77^.

### Genomic inflation

We observed a marked genomic inflation of the test statistics even after controlling for population stratification (linear mixed model) arising mainly from common markers; λ_GC_ in the primary meta-analysis (combined ancestries and combined sexes) was 1.06 and 1.37 for all and only common coding and splice site markers considered herein, respectively **(Supplementary Figures 3, 7** and **13, Supplementary Table 16)**. Such inflation is expected for a highly polygenic trait like WHRadjBMI, for studies using a non-random set of variants across the genome, and is consistent with our very large sample size^79,82,83^

### Conditional analyses

The RAREMETAL R-package^77^ was used to identify independent WHRadjBMI association signals across all ancestries and European meta-analysis results. RAREMETAL performs conditional analyses by using covariance matrices to distinguish true signals from the shadows of adjacent significant variants in LD. First, we identified the lead variants (P<2×10^−7^) based on a 1Mb window centered on the most significantly associated variant. We then conditioned on the lead variants in RAREMETAL and kept new lead signals at P<2×10^−7^ for conditioning in a second round of analysis. The process was repeated until no additional signal emerged below the pre-specified P-value threshold (P<2×10^−7^).

To test if the associations detected were independent of the previously published WHRadjBMI variants^10,14,16^, we performed conditional analyses in the stage 1 discovery set if the GWAS variant or its proxy (r^2^≥0.8) was present on the ExomeChip using RAREMETAL^77^. All variants identified in our meta-analysis and the previously published variants were also present in the UK Biobank dataset^84^. This dataset was used as a replacement dataset if a good proxy was not present on the ExomeChip as well as a replication dataset for the variants present on the ExomeChip. All conditional analyses in the UK Biobank dataset were performed using SNPTEST^85–87^. The conditional analyses were carried out reciprocally, conditioning on the ExomeChip variant and then the previously published variant. An association was considered independent of the previously published association if there was a statistically significant association detected prior to the conditional analysis (P<2×10^−7^) with both the exome chip variant and the previously published variant, and the observed association with both or either of the variants disappeared upon conditional analysis (P>0.05). A conditional p-value between 9×10^−6^ and 0.05 was considered inconclusive. However, a conditional p-value < 9×10^−6^ was also considered suggestive.

### Stage 2 meta-analyses

In our Stage 2, we sought to validate a total of 70 variants from Stage 1 that met P<2×10^−6^ in two independent studies, the UK Biobank (Release 1^84^) and Iceland (deCODE), comprising 119,572 and 12,605 individuals, respectively (Supplementary Tables 1-3). The same QC and analytical methodology were used for these studies. Genotyping, study descriptions and phenotype descriptives are provided in **Supplementary Tables 1-3**. For the combined analysis of Stage 1 plus 2, we used the inverse-variance weighted fixed effects meta-analysis method. Significant associations were defined as those nominally significant (P<0.05) in the Stage 2 study and for the combined meta-analysis (Stage 1 plus Stage 2) significance was set at P<2×10^−7^ (0.05/~250,000 variants).

### Pathway enrichment analyses: EC-DEPICT

We adapted DEPICT, a gene set enrichment analysis method for GWAS data, for use with the ExomeChip (‘EC-DEPICT’); this method is also described in a companion manuscript^22^. DEPICT’s primary innovation is the use of “reconstituted” gene sets, where many different types of gene sets (e.g. canonical pathways, protein-protein interaction networks, and mouse phenotypes) were extended through the use of large-scale microarray data (see Pers et al.^21^ for details). EC-DEPICT computes p-values based on Swedish ExomeChip data (Malmö Diet and Cancer (MDC), All New Diabetics in Scania (ANDIS), and Scania Diabetes Registry (SDR) cohorts, N=ll,899) and, unlike DEPICT, takes as input only the genes directly containing the significant (coding) variants rather than all genes within a specified amount of linkage disequilibrium (see **Supplementary Note 2).**

Two analyses were performed for WHRadjBMI ExomeChip: one with all variants p<5×l0^−4^ significant gene sets in 25 meta-gene sets, FDR <0.05) and one with all variants>1 Mb from known GWAS loci^10^ (26 significant gene sets in 13 meta-gene sets, FDR <0.05). Affinity propagation clustering^88^ was used to group highly correlated gene sets into “meta-gene sets”; for each meta-gene set, the member gene set with the best p-value was used as representative for purposes of visualization (see Supplementary Note). DEPICT for ExomeChip was written using the Python programming language, and the code can be found at https://github.com/RebeccaFine/obesity-ec-depict.

### Pathway enrichment analyses: PASCAL

We also applied the PASCAL pathway analysis tool^23^ to exome-wide association summary statistics from Stage 1 for all coding variants. The method derives gene-based scores (both SUM and MAX statistics) and subsequently tests for over-representation of high gene scores in predefined biological pathways. We used standard pathway libraries from KEGG, REACTOME and BIOCARTA, and also added dichotomized (Z-score>3) reconstituted gene sets from DEPICT^21^. To accurately estimate SNP-by-SNP correlations even for rare variants, we used the UK10 K data (TwinsUK^89^ and ALSPAC^90^ studies, N=3781). In order to separate the contribution of regulatory variants from the coding variants, we also applied PASCAL to association summary statistics of only regulatory variants (20 kb upstream) and regulatory+coding variants from the Shungin et al^10^ study. In this way, we could comment on what is gained by analyzing coding variants available on ExomeChip arrays. We performed both MAX and SUM estimations for pathway enrichment. MAX is more sensitive to genesets driven primarily by a single signal, while SUM is better when there are multiple variant associations in the same gene.

### Monogenic obesity enrichment analyses

We compiled two lists consisting of 31 genes with strong evidence that disruption causes monogenic forms of insulin resistance or diabetes; and 8 genes with evidence that disruption causes monogenic forms of lipodystrophy. To test for enrichment of association, we conducted simulations by matching each gene with others based on gene length and number of variants tested, to create a matched set of genes. We generated 1,000 matched gene sets from our data, and assessed how often the number of variants exceeding set significance thresholds was greater than in our monogenic obesity gene set.

### Variance explained

We estimated the phenotypic variance explained by the association signals in Stage 1 all ancestries analyses for men, women, and combined sexes^91^. For each associated region, we pruned subsets of SNPs within 500 kb, as this threshold was comparable with previous studies, of the SNPs with the lowest P-value and used varying P value thresholds (ranging from 2×l0^−7^ to 0.02) from the combined sexes results. Additionally, we examined all variants and independent variants across a range of MAF thresholds. The variance explained by each subset of SNPs in each strata was estimated by summing the variance explained by the individual top coding variants. For the comparison of variance explained between men and women, we tested for the significance of the differences assuming that the weighted sum of chi-squared distributed variables tend to a Gaussian distribution ensured by Lyapunov’s central limit theorem.^91,92^

### Cross-trait lookups

To carefully explore the relationship between WHRadjBMI and related cardiometabolic, anthropometric, and reproductive traits, association results for the 51 WHRadjBMI coding SNPs were requested from existing or on-going meta-analyses from 7 consortia, including ExomeChip data from GIANT (BMI, height), Global Lipids Genetics Consortium Results (GLGC) (total cholesterol, triglycerides, HDL-cholesterol, LDL-cholesterol), International Consortium for Blood Pressure (IBPC)^93^ (systolic and diastolic blood pressure), Meta-Analyses of Glucose and Insulin-related traits Consortium (MAGIC) (glycemic traits), and DIAbetes Genetics Replication And Meta-analysis (DIAGRAM) consortium (type 2 diabetes).).^22,29–25^ For coronary artery disease, we accessed 1000 Genomes Project-imputed GWAS data released by CARDI0GRAMplusC4D^94^ and for the ReproGen consortium (age at menarche and menopause) we used a combination of ExomeChip and 1000 Genomes Project-Imputed GWAS data. Heatmaps were generated in R v3.3.2 using gplots (https://CRAN.R-project.org/package=gplots). We used Euclidean distance based on p-value and direction of effect and complete linkage clustering for the dendrograms.

### GWAS Catalog Lookups

In order to determine if significant coding variants were associated with any related cardiometabolic and anthropometric traits, we also searched the NHGRI-EBI GWAS Catalog for previous variant-trait associations near our lead SNPs (+/− 500 kb). We used PLINK to calculate LD for variants using ARIC study European participants. All SNVs within the specified regions with an r^2^ value>0.7 were retained from NHGRI-EBI GWAS Catalog for further evaluation^37^. Consistent direction of effect was based on WHR-increasing allele, LD, and allele frequency. Therefore, when a GWAS Catalog variant was not identical or in high LD (r^2^>0.9) with the WHR variant, and MAF >0.45, we do not comment on direction of effect.

### Body-fat percentage associations

We performed body fat percent and truncal fat percent look-up of 48 of the 56 identified variants (tables 1 and 2) that were available in the UK Biobank, Release l^84^, data (notably some of the rare variants in table 1 and 2 were not available) to further characterize their effects on WHRadjBMI. Genome-wide association analyses for body fat percent and truncal fat percent were carried out in the UK Biobank. Prior to analysis, phenotype data were filtered to exclude pregnant or possibly pregnant women, individuals with body mass index<15, and without genetically confirmed European ancestry, resulting in a sample size of 120,286. Estimated measures of body fat percent and truncal fat percent were obtained using the Tanita BC418MA body composition analyzer (Tanita, Tokyo, Japan). Individuals were not required to fast and did not follow any specific instructions prior to the bioimpedance measurements. SNPTEST was used to perform the analyses based on residuals adjusted for age, 15 principle components, assessment center and the genotyping chip^85^.

### Collider bias

In order to evaluate SNPs for possible collider bias^18^, we used results from a recent association analysis from GIANT on BMI^25^. For each significant SNP identified in our additive models, WHRadjBMI associations were corrected for potential bias due to associations between each variant and BMI (See **Supplementary Note 1** for additional details). Variants were considered robust against collider bias if they met Bonferroni-corrected significance following correction (P_corrected_<9.09×l0^−4^, 0.05/55 variants examined).

### Drosophila RNAi knockdown experiments

For each gene in which coding variants were associated with WHRadjBMI in the final combined meta-analysis (P<2×l0^−7^), its corresponding Drosophila orthologues were identified in the Ensembl ortholog database (www.ensembl.org), when available. Drosophila triglyceride content values were mined from a publicly available genome-wide fat screen data set^45^ to identify potential genes for follow-up knockdowns. Estimated values represent fractional changes in triglyceride content in adult male flies. Data are from male progeny resulting from crosses of male UAS-RNAi flies from the Vienna Drosophila Resource Center (VDRC) and Hsp70-GAL4; Tub-GAL8ts virgin females. Two-to-five-day-old males were sorted into groups of 20 and subjected to two one-hour wet heatshocks four days apart. On the seventh day, flies were picked in groups of eight, manually crushed and sonicated, and the lysates heat-inactivated for 10 min in a thermocycler at 95 °C. Centrifuge-cleared supernatants were then used for triglyceride (GPO Trinder, Sigma) and protein (Pierce) determination. Triglyceride values from these adult-induced ubiquitous RNAi knockdown individuals were normalized to those obtained in parallel from non-heatshocked progeny from the very same crosses. The screen comprised one to three biological replicates. We followed up each gene with a >0.2 increase or >0.4 decrease in triglyceride content.

Orthologues for two genes were brought forward for follow-up, *DNAH10* and *PLXND1*. For both genes, we generated adipose tissue (cg-Gal4) and neuronal (elav-Gal4) specific RNAi-knockdown crosses to knockdown transcripts in a tissue specific manner, leveraging upstream activation sequence (UAS)-inducible short-hairpin knockdown lines, available through the VDRC (Vienna *Drosophila* Resource Center). Specifically, elav-Gal4, which drives expression of the RNAi construct in post mitotic neurons starting at embryonic stages all the way to adulthood, was used. Cg drives expression in the fat body and hemocytes starting at embryonic stage 12, all the way to adulthood. We crossed male UAS-RNAi flies and elav-GAL4 or CG-GAL4 virgin female flies. All fly experiments were carried out at 25°C. Five-to-seven-day-old males were sorted into groups of *20*, weighed and homogenated in PBS with 0.05% Tween with Lysing Matrix D in a beadshaker. The homogenate was heat-inactivated for 10 min in a thermocycler at 70°C. 10μl of the homogenate was subsequently used in a triglyceride assay (Sigma, Serum Triglyceride Determination Kit) which was carried out in duplicate according to protocol, with one alteration: the samples were cleared of residual particulate debris by centrifugation before absorbance reading. Resulting triglyceride values were normalized to fly weight and larval/population density. We used the non-parametric Kruskall-Wallis test to compare wild type with knockdown lines.

### Expression quantitative trait loci (eQTLs) analysis

We queried the significant variant (Exome coding SNPs)-gene pairs associated with eGenes across five metabolically relevant tissues (skeletal muscle, subcutaneous adipose, visceral adipose, liver and pancreas) with at least 70 samples in the GTEx database^46^. For each tissue, variants were selected based on the following thresholds: the minor allele was observed in at least 10 samples, and the minor allele frequency was≥0.01. eGenes, genes with a significant eQTL, are defined on a false discovery rate (FDR)^95^ threshold of ≤0.05 of beta distribution-adjusted empirical p-value from FastQTL. Nominal p-values were generated for each variant-gene pair by testing the alternative hypothesis that the slope of a linear regression model between genotype and expression deviates from 0. To identify the list of all significant variant-gene pairs associated with eGenes, a genome-wide empirical p-value threshold^64^, pt, was defined as the empirical p-value of the gene closest to the 0.05 FDR threshold, pt was then used to calculate a nominal p-value threshold for each gene based on the beta distribution model (from FastQTL) of the minimum p-value distribution f(pmin) obtained from the permutations for the gene. For each gene, variants with a nominal p-value below the gene-level threshold were considered significant and included in the final list of variant-gene pairs^64^. For each eGene, we also listed the most significantly associated variants (eSNP). Only these exome SNPs with r^2^>0.8 with eSNPs were considered for the biological interpretation (Supplementary eQTL GTEx).

We also performed cis-eQTL analysis in 770 METSIM subcutaneous adipose tissue samples as described in Civelek, et al.^96^ A false discovery rate (FDR) was calculated using all p-values from the cis-eQTL detection in the q-value package in R. Variants associated with nearby genes at an FDR less than 1% were considered to be significant (equivalent p-value<2.46 χ 10^−4^).

For loci with more than one microarray probeset of the same gene associated with the exome variant, we selected the probeset that provided the strongest LD r2 between the exome variant and the eSNP. In reciprocal conditional analysis, we conditioned on the lead exome variant by including it as a covariate in the cis-eQTL detection and reporting the p-value of the eSNP and vice versa. We considered the signals to be coincident if both the lead exome variant and the eSNP were no longer significant after conditioning on the other and the variants were in high pairwise LD (r2>0.80).

For loci that also harbored reported GWAS variants, we performed reciprocal conditional analysis between the GWAS lead variant and the lead eSNP. For loci with more than one reported GWAS variant, the GWAS lead variant with the strongest LD r2 with the lead eSNP was reported.

### Penetrance analysis

Phenotype and genotype data from the UK Biobank (UKBB) were used for the penetrance analysis. Three of 16 rare and low frequency variants (MAF≤1%) detected in the final Stage 1 plus 2 meta-analysis were available in the UKBB and had relatively larger effect sizes (>0.90). The phenotype data for these three variants were stratified with respect to waist-to-hip ratio (WHR) using the World Health Organization (WHO) guidelines. These guidelines consider women and men with WHR greater than 0.85 and 0.90 as obese, respectively. Genotype and allele counts were obtained for the available variants and these were used to calculate the number of carriers of the minor allele. The number of carriers for women, men and all combined was then compared between two strata (obese vs. non-obese) using a χ2 test. The significance threshold was determined by using a Bonferroni correction for the number of tests performed (0.05/9=5.5×l0^−3^)).

## DATA AVAILABILITY

Summary statistics of all analyses are available at https://www.broadinstitute.org/collaboration/giant/.

